# Emergence of neocortical function in heterotopic neurons

**DOI:** 10.1101/2024.01.17.576031

**Authors:** Sergi Roig-Puiggros, Maëlle Guyoton, Dmitrii Suchkov, Aurélien Fortoul, Sabine Fièvre, Giulio Matteucci, Emma Maino, Charlie G. Foucher, Daniel Fuciec, Esther Klingler, Fiona Francis, Marat Minlebaev, Sami El-Boustani, Françoise Watrin, Jean-Bernard Manent, Denis Jabaudon

## Abstract

Brains come in various sizes and shapes, yet how neuronal position constrains the type of circuits that they can form remains largely unknown. The spatial layout of anatomical structures with corresponding functions varies widely across species (*J*-*4*). Also, during evolution, anatomical structures have duplicated and then diverged to generate new circuits and functions (*5*, *6*). Thus, it is critical to understand how the position of neurons constrains their integration into circuits and, ultimately, their function. To address this question, we studied *EmlJ* knockout mice in which subsets of neocortical neurons form a new structure below the neocortex termed heterotopia (Ht). We examined how this new location affects the molecular identity, topography, input-output circuit connectivity, electrophysiology, and functional properties of these neurons. Our results reveal a striking conservation of the cellular features and circuit properties of Ht neurons, despite their abnormal location and misorientation. Supporting this observation, these neurons were able to functionally substitute for overlying neocortical neurons in a behaviorally relevant task when the latter were optogenetically silenced. Hence, specific neuronal identities and associated function can be reproduced in altered anatomical settings, revealing a remarkable level of self-organization and adaptability of neocortical circuits.

## Main text

To address whether neurons and the circuits they form need to settle in their appropriate locations to functionally engage, we studied telencephalon-specific Eml1 knockout mice (*Emx1:Cre; Eml1^lox/lox^* or cKO). In these mice, neocortical apical progenitors detach from the ventricular wall during embryogenesis, leading to secondary ectopic neuronal generation with the formation of a large subcortical heterotopia (Ht) (*7–10*) (Fig. 1A and fig. S1). We sequentially examined the molecular and spatial identity of Ht neurons, their output connectivity, electrophysiological features, input connectivity, and functional relevance.

**Figure 1.**
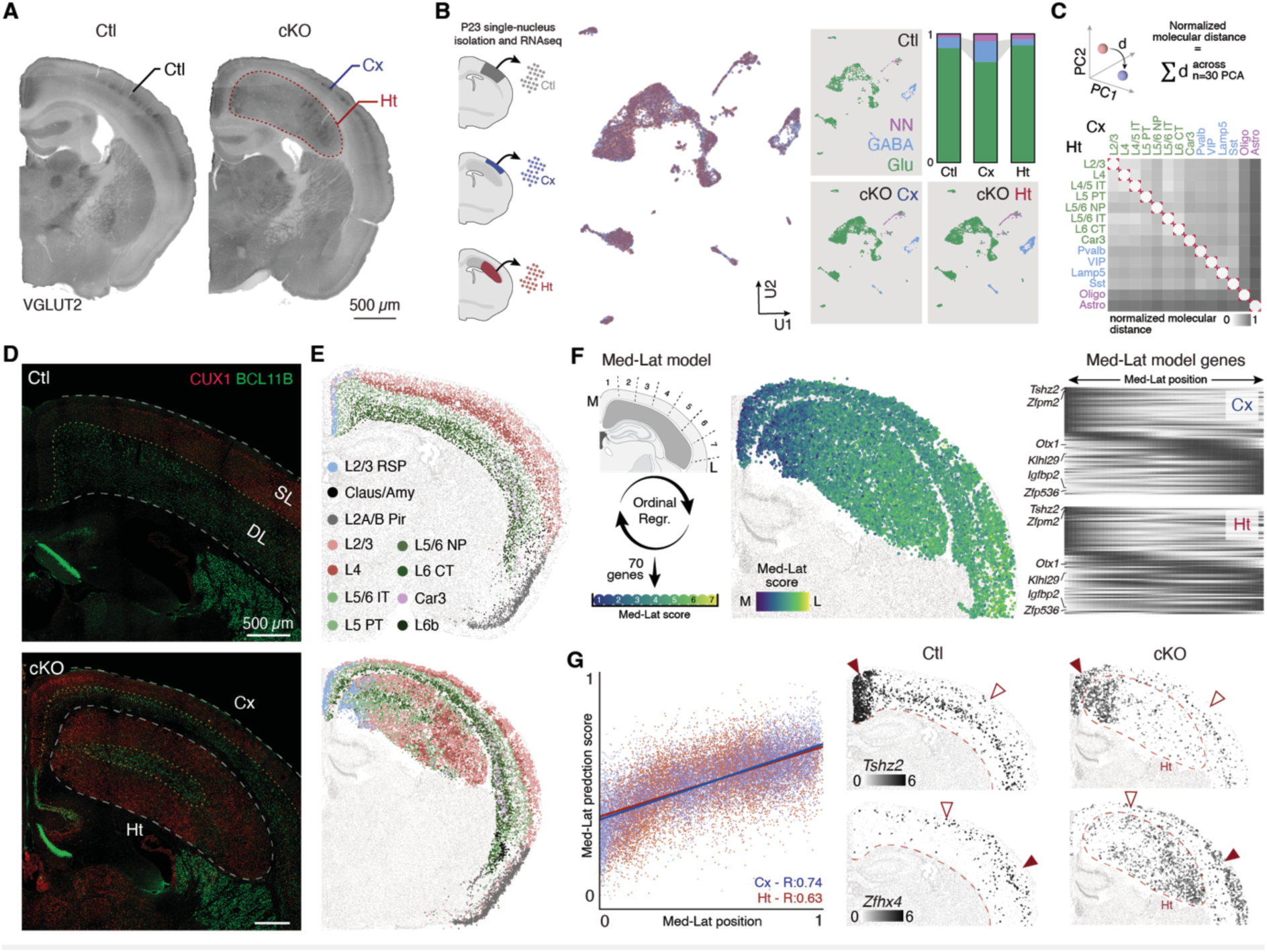
Ht neurons reproduce the molecular and spatial identities of cortical neurons. **(A)** Coronal section of P7 Ctl (left) and cKO (right) brains. **(B)** Left: Schematic representation of the single-nucleus (sn) isolation and RNA sequencing strategy. Middle: UMAP (Ul and U2) representation of snRNAseq cells color-coded by sample. Right: UMAP representation of snRNAseq datasets split by sample and color-coded by cell-classes: glutamatergic neurons (green), GABAergic neurons (blue) and non-neuronal cells (purple) and their relative proportions. **(C)** Top: Schematic representation of the molecular distance calculation (based on the 30 first principal components) between Ht and Cx cell types. Bottom: Heatmap of PC distances between Ht and Cx cell types. **(D)** Coronal section of an adult Ctl (top) and cKO (bottom) brain stained for CUXl and BCLllB. **(E)** Spatial transcriptomics dataset of a wild-type coronal section (*15*)(top) and *Ht* brain coronal section (bottom) color-coded by cortical neuronal type. **(F)** Left: Ordinal regression model of neuronal position based on 7 cortical medio-lateral (med-lat) bins. Middle: Med-lat position score of cortical and Ht neurons. Right: Spatial distribution of the genes according to their med-lat score. **(G)** Left: Correlation between actual med-lat position (x) and predicted med-lat score (y) in the cortex (blue) and Ht (red). Illustrative distribution of medially-(*e.g. Tshz2*) and laterally-(*e.g. Zfhx4*) enriched genes in the cortex and Ht. Filled and empty arrowheads indicate high and low expression levels, respectively. RNAseq: RNA sequencing, UMAP: Uniform manifold approximation projection, PCA: principal component analysis, PC: principal component, L: layer, RSP: retrosplenial, Claus/Amy: claustrum/amygdala, Pir: piriform, NP: near-projecting, CT: cortico-thalamic, IT: intra-telencephalic, PT: pyramidal tract, Med-Lat: mediolateral, M: medial, L: lateral, Ordinal regr: ordinal regression model.

### Molecular and spatial identity

To assess which cell types are present in the Ht, we performed single-nucleus RNA sequencing of the Ht, the overlying neocortex (“cortex”, Cx), and of control cortex from wild-type littermates (*Eml1^lox/lox^*; Ctl). To unbiasedly predict cortical cell identity, we used a feedforward neural network trained on a primary somatosensory (SSp) cortex single-cell RNA sequencing dataset (see Methods). We found that cellular diversity in the Ht was comparable to that of the cortex, as indicated by overlapping distributions of cells following UMAP dimensionality reduction (Fig. 1B, fig. S2A-D). The proportion of deep-layer neurons was slightly decreased in the Ht (fig. S2E), possibly because, when cKO apical progenitors first detach from the ventricular wall, subsets of deep-layer neurons have already been generated (*7*). The proportion of GABAergic interneurons was also decreased, suggesting non-cell autonomous impaired migration into the Ht from the subpallium, where GABAergic neurons are generated (Fig. 1B, fig. S2E, S3B) (*11*). Despite these differences in abundance, across cell types, the molecular identity of Ht cells could not be distinguished from that of correctly positioned cortical cells, as demonstrated by measuring the molecular distances between corresponding cell types in Ht, Cx, and Ctl (Fig. 1C, fig. S2F-G). Together, these results indicate that despite their abnormal location, the diversity and molecular identities of Ht cells largely reproduce those of correctly positioned cortical cells.

In the cortex, neurons are organized radially into layers, and medio-laterally into areas.

To assess the radial organization of Ht neurons, we first performed immunostainings for BCLIIB (expressed by deep-layer cortical neurons) and CUXI (expressed by superficial-layer neurons) (*12*). We observed that, although BCLIIB- and CUXI-expressing neurons were spatially organized within the Ht, their layout was not laminar. Instead, they organized in a concentric pattern, in which "deep-layer", BCLIIB-expressing neurons were found in the "core" of the Ht, while "superficial-layer", CUXI-expressing neurons occupied the "shell" of the Ht (Fig. ID, fig. S3A). Using immunostaining for ANK3, a protein localized at the axonal initial segment, we discovered that in contrast to cortical pyramidal neurons, which are mostly radially oriented, Ht neurons have random orientations (fig. S3C). Yet, despite this cytoarchitectural disorganization, expression of BCLIIB and CUXI remained essentially mutually exclusive, as is the case in the cortex. This suggests that cellular orientation does not affect molecular identities (fig. S3A). Core neurons were born before shell neurons, as demonstrated using EdU birthdating (fig. S3D), which recapitulates the normal sequence of cortical neuron birth, in which deep-layer neurons are born earlier than superficial layer neurons. Hence the sequential developmental timing of distinct neuron types is reproduced in the Ht.

We next used spatial transcriptomics to assess the medio-lateral organization of molecularly defined classes of Ht cells (Fig. IE-G, fig. S4). Using combinations of genes to define specific cortical neuron types, we found that the medio-lateral distribution of excitatory Ht neurons types matched with that of cortical neurons (control dataset from (*13*)). For example, retrosplenial-type neurons were present medially, as is normally the case in the cortex (Fig. 1E). To unbiasedly assess the tangential distribution of Ht neurons, we built an ordinal regression model that identified seventy core genes predicting the medio-lateral position of neocortical neurons (Fig. 1F, fig. S5). The latter predicted cell position with the same accuracy for Ht neurons as for cortical neurons (Fig. 1G left). Tllustrating these results, transcripts such as *Tshz2* and *Zfhx4*, which are enriched in the medial and lateral cortex, respectively, had a corresponding distribution in the Ht (Fig. 1G right). Thus, the topographical medio-lateral molecular organization of Ht neurons reproduces that of cortical neurons.

### Output connectivity

Next, we examined whether the conserved molecular identity and topographical mapping of Ht neurons were matched by corresponding axonal projection patterns. To label and analyze efferent pathways, we first bulk-injected an anterograde adeno-associated viral vector (AAV) to express GFP into the Ht. We then performed 3D reconstructions of these pathways in cleared brains (fig. S6A). We observed two main classes of long-distance projections originating from the Ht. First, corticofugal-like type projections, which targeted subcerebral structures such as the thalamus, superior colliculus, hindbrain, and spinal cord. Second, intracortical-like type projections, which fanned out from the Ht in distinct fascicles towards the ipsilateral and contralateral hemisphere, including the contralateral Ht (fig. S6A-B).

To better characterize the topographical arrangement of intracortical-like-type projections, we first focused on interhemispheric projections. Callosal projection neurons normally send their axon to the contralateral hemisphere in a homotopic manner, i.e. laterally located neurons project laterally, and medially located neurons medially (Fig. 2A left) (*14*, *15*). Using double injections of retrograde tracers (cholera toxin subunit B) conjugated with two different dyes (Alexa 555 and 647) in the lateral and medial cortex, respectively, we found that this layout is reproduced in the Ht, consistent with a conserved topology of long-range intracortical-like projections (Fig. 2A right, fig. S6B).

**Figure 2.**
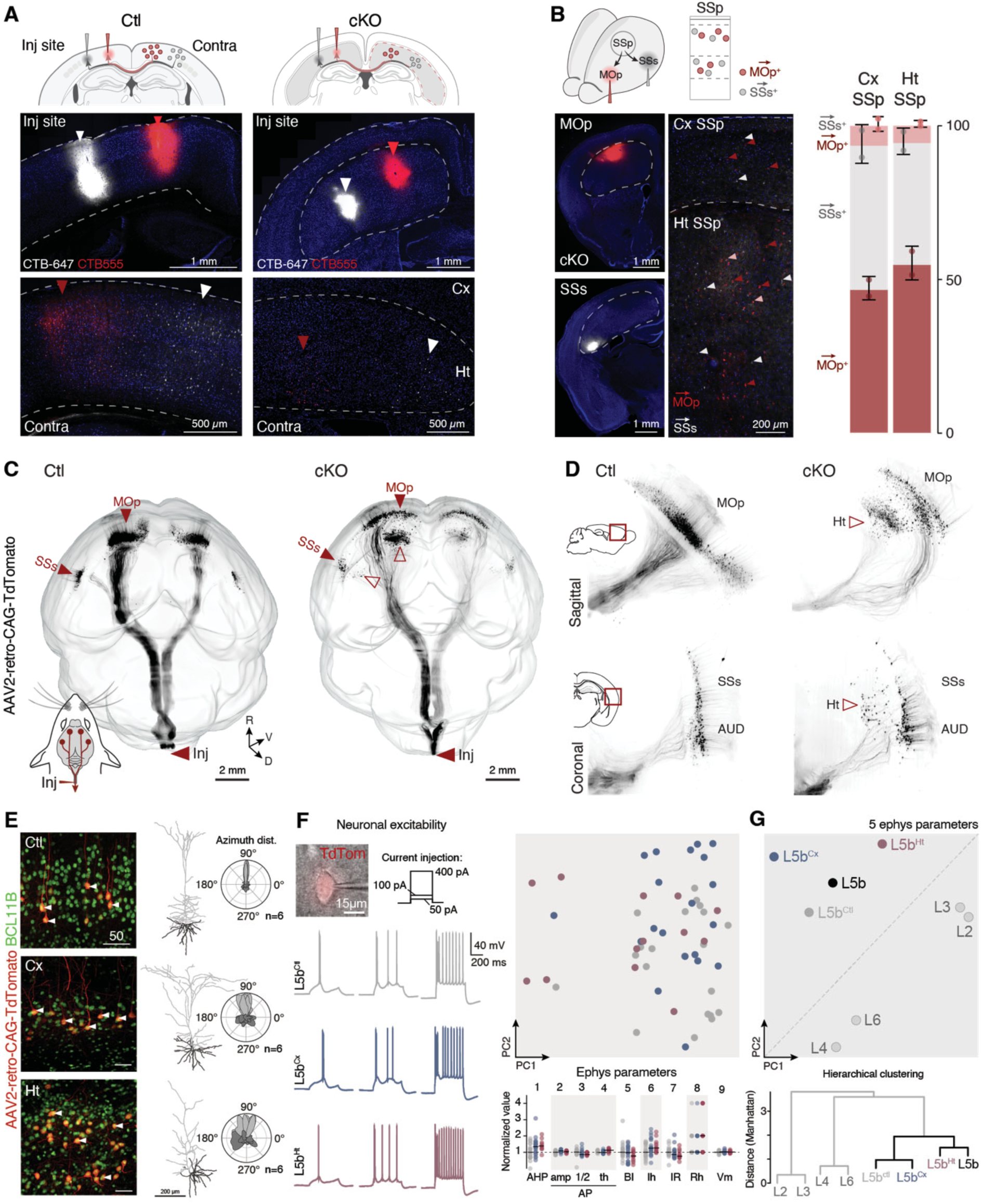
Ht neurons have cortical neuron-like output connectivity and electrophysiological properties. Top: Schematic representation of the homotopic callosal projections retrograde tracing strategy. Middle: Illustration of the CTB injection sites. Bottom: Medial (red) and lateral (white) CTB^+^ neurons, retrogradely traced from the contralateral cortex or Ht. (**B)** Top: Schematic representation of SSp ICPN retrograde tracing from MOp and SSs. Bottom left: Illustration of the CTB injection sites in MOp and SSs. Bottom right: Retrogradely-labeled MOp projecting (red) and SSs projecting (white) CTB^+^ neurons and their proportions in Cx SSp and Ht SSp. **(C)** Spinal-cord-projecting neurons in Ctl (left) and cKO (right) brains, retrogradely labeled by injection of a TdTomato-expressing AAV in the cervical cord (arrowhead indicates injection site). 3D-reconstructed cleared brains. **(D)** Digitally-reconstructed 2D sections at the level of MOp (sagittal) and SSs (coronal) for Ctl (left) and *Ht* (right) brains. **(E)** Left: Photomicrograph showing immunofluorescence against BCLllB in spinal-cord-projecting retrogradely-labeled TdTomato^+^ Ht neurons. Right: biocytin reconstruction of CST projecting neurons (apical dendrite: lightgrey, basal dendrite: darkgrey) and their respective azimuth density distribution **(F)** Patch-clamp recordings on TdTomato^+^ spinal-cord projecting neurons (i.e. L5b-type neurons) from Ctl and cKO brains. Left: sample recordings of the response to depolarization steps from Ctl, Cx and Ht. Right: Principal Component Analysis (PCA) of the 9 recorded parameters for all the sampled neurons (Ctl =2l, *Cx*=20, *Ht*=l3). Bottom: summary of recording results for 9 electrophysiological parameters. **(G)** Top: PCA analysis of 5 electrophysiological parameters for specific neuron types (see Methods). Bottom: Hierarchical clustering of the same dataset. CTB: cholera toxin subunit B, ICPN: inter-areal cortical projecting neuron, Contra: contralateral, Ht: heterotopia, Cx: cortex, SSp: primary somatosensory cortex, MOp: primary motor cortex, SSs: secondary somatosensory cortex, CST: cortico-spinal tract, Inj: injection site, AUD: primary auditory cortex, L5b: layer 5b neurons, ampl: amplitude, AP: action potential, pA: picoamp, mV: millivolt, ms: millisecond, th: threshold, Vm: membrane voltage, R: resistance, PC: principal component.

In the primary somatosensory cortex (SSp), intracortically-projecting neurons have mutually exclusive projections to either the secondary somatosensory cortex (SSs) or to the primary motor cortex (MOp) (*16*-*19*). This can be demonstrated by simultaneous retrograde labeling from SSs and MOp and assessing the proportion of single-labeled somas - around 90% in Ctl mice (fig. S6C). Given that molecular topographical mapping in the Ht matches that of the Cx, we hypothesized that Ht regions corresponding to SSp, SSs and MOp would be present and reproduce the connectivity present in the cortex. Confirming this possibility, double retrograde labeling from the region of the Ht located below SSs and below MOp (Fig. 2B left) revealed largely mutually exclusively labeled neuronal somas in the region below SSp. This, indicates that cell-type specific intracortical-like connectivity is present in the Ht (Fig. 2B).

The corticospinal tract is a key motor pathway that projects from the cortex to the spinal cord. To assess the connectivity of Ht neurons with the spinal cord, we performed labeling from the cervical spinal cord by injecting a retrogradely-transported AAV expressing TdTomato. Retrogradely-labeled somas in the Ht were found in areas immediately below the MOp and SSs, where most corticospinal neurons are normally found (Fig. 2C, D). Hence, the molecular topographical organization of the Ht is accompanied by a corresponding topographical organization of the connectivity of the neurons that compose it. Ht spinal cord-projecting neurons express BCLIIB, as corticospinal neurons normally do (*20*), supporting a conserved correspondence between molecular identity and connectivity in Ht neurons (Fig. 2D left, fig. S2, S4).

To assess the morphology of retrogradely-labeled Ht neurons, we filled them with biocytin during whole-cell patch-clamp recordings. We found that they polarized appropriately into a pyramidal shape (Fig. 2D right, fig. S6D), although they were abnormally oriented (fig. S3C). Remarkably, despite these abnormal orientations, they were still able to project to the spinal cord.

### Electrophysiology

To assess whether electrophysiological features of Ht neurons match corresponding populations of cortical neurons, we performed whole-cell patch-clamp recordings of the retrogradely labeled spinal cord-projecting neurons. Based on the measurement of nine electrophysiological parameters, Ht neurons were indistinguishable from correctly positioned cortical neurons, including in terms of resting membrane potential, excitability, and action potential firing properties (Fig. 2E, fig. S6E-G).

The distinct types of glutamatergic neurons of the cortex can, to some extent, be distinguished by their electrophysiological features (*21*). To address cell-type specific electrophysiological properties of spinal cord-projecting Ht neurons, we took advantage of a dataset reporting the distinct properties of L5B (i.e. corticospinal neurons), L6, L4, and L2/3 excitatory neurons (*21*). Using principal component analysis of five electrophysiological features and hierarchical clustering of these neuronal types, we found that the electrophysiological properties of Ht spinal cord-projecting neurons were similar to those of genuine corticospinal neurons, and distinct from those of other neuron types (Fig. 2F). Therefore, not only are overall cortical electrophysiological properties reproduced in Ht neurons, but the specific components of the electrophysiological signatures that characterize this cell type are also maintained.

### Somatosensory input

We then examined whether Ht neurons receive input from the thalamus as cortical neurons normally do. To address this question, we first performed anterograde labeling from the ventroposterior medial nucleus (VPM) of the thalamus, which projects to the barrel field of SSp (SSp-bfd) (*22*). This revealed that thalamic axons invade the lateral part of the Ht, immediately across from SSp-bfd (Fig. 3A, fig. S7A). In the posteromedial part of SSp, thalamic afferents are somatotopically organized, mirroring the layout of the whisker pad. During the early postnatal period, L4 neurons form anatomical clusters called barrels, with individual barrels receiving input predominantly from a single whisker (*23*). Consistent with somatotopically organized thalamic afferents within the Ht, VGLUT2 labeling of presynaptic thalamic axons revealed a topographical mapping of whiskers into barrels comparable to that found in the overlying cortex - yet somewhat distorted compared to control cortex and in line with previous results (Fig. 3B, fig. S7B, (*24*)). Ht and cortical neurons thus have corresponding thalamic input-driven topologies. Ht neurons clustered to form barrel-like structures around VPM thalamic afferents. These neurons expressed RORB, corresponding to a L4-type identity in the cortex (Fig. 3C). Hence, in agreement with our findings on output connectivity (Fig. 2A-D), Ht neurons display a conserved correspondence between molecular identity and input connectivity.

**Figure 3.**
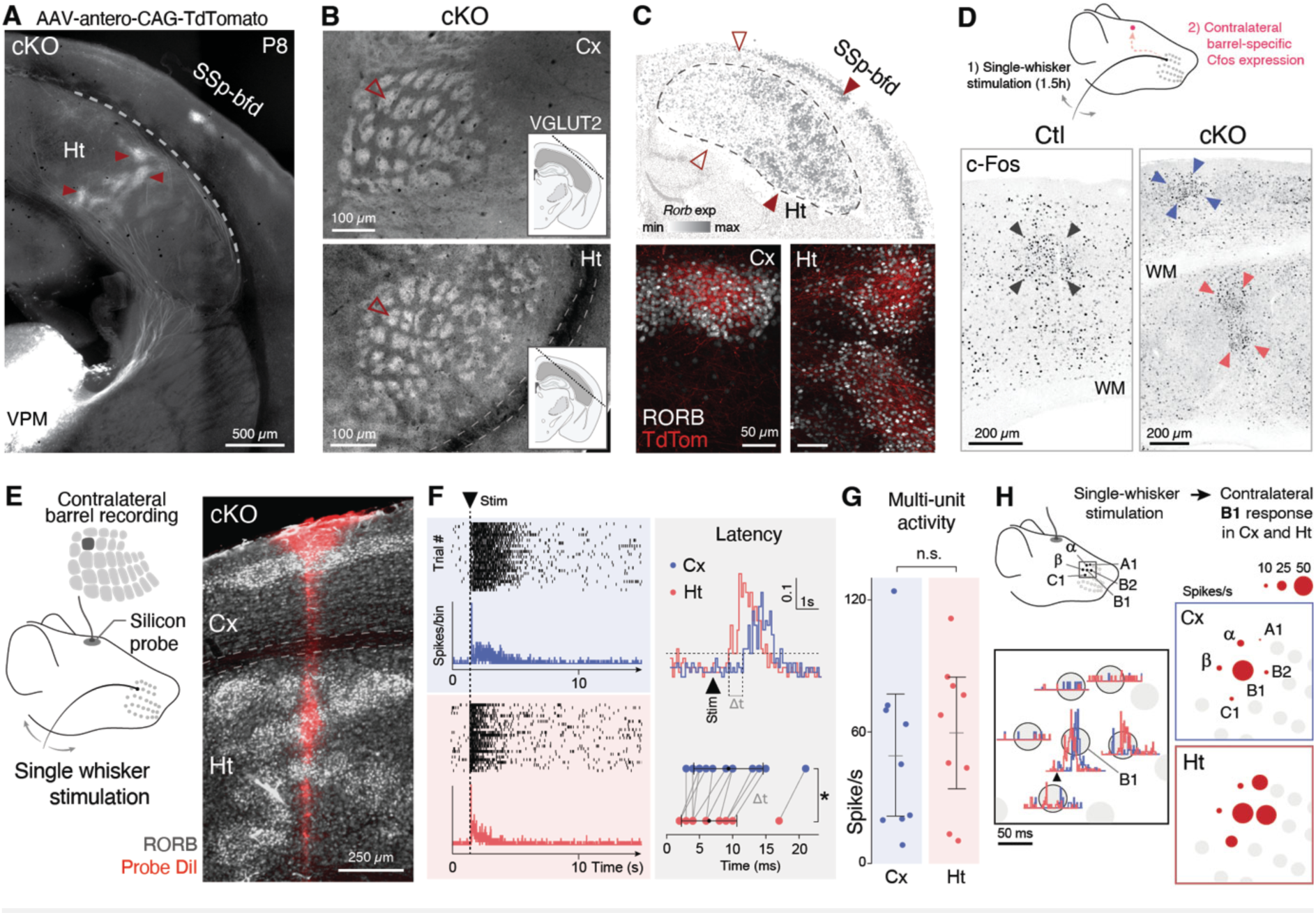
Somotosensory input to Ht neurons is anatomically and functionally conserved. **(A)** Anterograde tracing tracing of thalamic VPM afferents to Cx and Ht. Arrowheads highlight axonal arborization into barrel-like structures. **(B)** VGLUT2 labeling of thalamic terminals on flattened preparations highlights the somatotopic organization of Ht input. Empty arrowhead points at the Bl barrel and its Ht equivalent. **(C)** Top: MERFISH spatial distribution of RORB-expressing cells (i.e. L4-like). white-arrow: low expression, red-arrow: high expression. Bottom: Thalamic afferents (high-magnification from (A)) cluster around RORB-expressing cells in the Cx (left) and Ht (right). **(D)** Top: Schematic illustration of the single-whisker stimulation paradigm. Bottom: c-Fos-expressing neurons are found in a single cortical barrel and underlying Ht. Arrowheads highlight barrel and Ht barrel-equivalent locations. **(E)** Left: Schematic representation barrel responses recording (multisite silicon probe) upon single whisker stimulation. Right: illustrative photomicrograph of a coronal section stained against RORB (i.e. L4 neurons) in which the probe position has been marked by Dil. **(F)** Left: Illustrative responses to single whisker stimulation in Cx (top, blue) and Ht (bottom, red). Right: Sample evoked responses (top) and average evoked response latencies (bottom, two-sided Wilcoxon rank sum test, n=l3). **(G)** Multi-unit activity in response to stimulation - a measure of sensitivity to input - in Cx and Ht (two-sided Wilcoxon rank sum test, n=9). **(H)** Left: Schematic representation of the experiment. Right: responses obtained in the Blwhisker representation recording upon stimulation of distinct whiskers (i.e. response tuning, n=l0). *: P < 0.05, SSp-bfd: primary somatosensory cortex barrel field, VPM: ventral posteromedial nucleus of the thalamus, Stim: stimulation, WM: white-matter.

To examine whether whisker-to-Ht connectivity was functional, we allowed mice to roam freely in an enriched environment and assessed neuronal activation through c-Fos expression. Prior to this, all but one whisker had been trimmed to allow for the exclusive stimulation of that whisker (*25*). Within the cortex, activation was confined to a single barrel, indicating that the Ht does not affect the normal functional topography between the whiskers and the cortex. In addition, however, in the Ht immediately below the activated cortical barrel, an activated barrel-like structure was also visible, revealing a functional and somatotopic duplication of barrel-to-forebrain pathways (Fig. 3D, fig. S7C).

To address the physiological dynamics of this connectivity, we simultaneously recorded from the cortex and the Ht with a multi-site silicon probe inserted orthogonally to the cortical surface (Fig. 3E; we used intrinsic optical imaging upon single-whisker stimulation to identify the electrode insertion site, see Methods). Recordings revealed phase-locked activation of neurons in both structures in response to single whisker deflections - Ht neurons were activated a few milliseconds before cortical neurons, consistent with a more proximal location with respect to thalamic input (Fig. 3F). Spiking rates were also similar between Ht neurons and cortical neurons, suggesting similar sensitivity to whisker deflections (Fig. 3G). Finally, the specificity of responses to single-whisker stimulations was preserved in Ht barrel equivalents - though less sharply than in the cortex - with the strongest evoked responses observed for the whisker corresponding to the recorded barrel equivalent (Fig. 3H). Together, these results reveal that cortical and Ht neurons together form functional units - "pseudo-columns" - which are somatotopically arranged and jointly respond to whiskers stimulation.

### Visual input

To examine whether Ht functional topographical mapping was limited to the somatosensory system or present for other sensory modalities, we measured the visual-evoked responses of Ht neurons using two-photon calcium imaging. We injected a GCAMP6f-expressing AAV (*26*) into the primary visual cortex (VTSp) and underlying Ht - prior to this, we had performed intrinsic optical signal imaging upon visual stimulation to identify the injection site (see Methods). We then placed an optical prism at this location, allowing us to simultaneously image evoked responses in cortical neurons and underlying Ht neurons in a single field of view (Fig. 4A).

**Figure 4.**
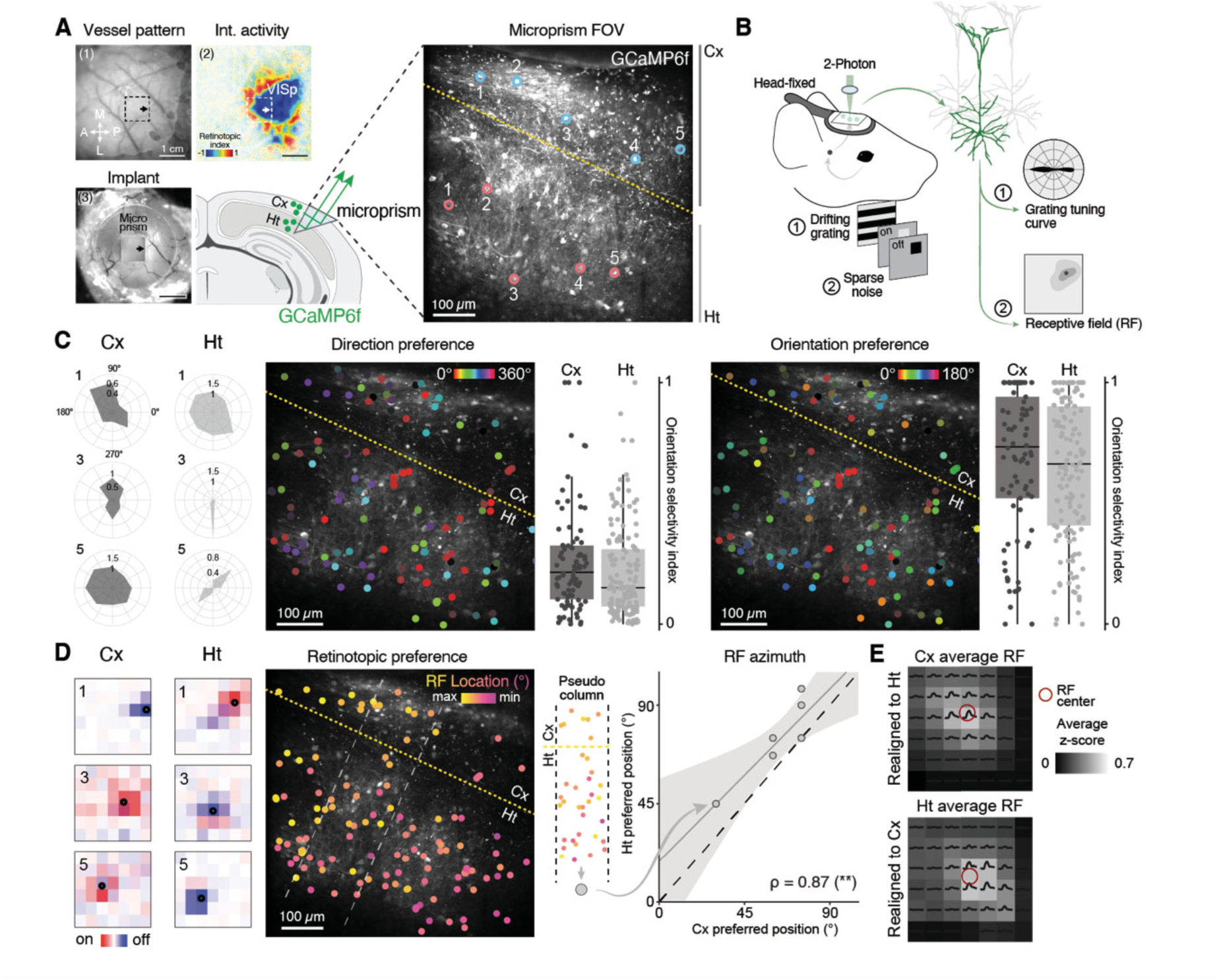
Retinotopic input to Ht neurons is anatomically and functionally conserved. **(A)** Left: Microprism implantation pipeline. Blood vessels serve as landmarks and intrinsic optical imaging allows identification of the injection site for GCaMP6f-encoding virus injection and implantation of a microprism across the Cx and the Ht. Neuronal GCaMP6f signals are imaged with a two-photon microscope. Right: The microprism allows visualization of GCaMP6f-expressing cortical (blue circles) and Ht (red circles) neurons in a single field of view. The dashed line represents the white matter. **(B)** Schematic representation of the visual stimulation paradigm, alternating drifting gratings and sparse noise. These stimulations are used to measure neurons direction and orientation tuning curves and receptive field, respectively. **(C)** Left: Polar plot of single-neuron direction/orientation preferences for select Cx and Ht neurons. Numbers correspond to the neurons highlighted in A. Middle: Spatial distribution of preferred direction, and its quantification. Right: Spatial distribution of preferred orientation, and its quantification (unpaired two-sided Wilcoxon test, direction: *P* = 0.l6, orientation: *P* = 0.034.**(D)** Left: heatmap representation of single-neuron receptive fields for select Cx and Ht neurons. Red: ON field, blue: OFF field. Same neurons as in C. Middle: Spatial distribution of retinotopic preference. Right: grouping neurons into spatially-confined "pseudocolumns" identifies corresponding preferences for receptive field azimuth in Cx and Ht neurons (Pearson coefficient of correlation (ρ), azimuth: P = 0.0l). **(E)** Cx and Ht neurons average receptive field recentered based on Ht or Cx respective receptive field location (n = 72 in Cx and n = l27 in Ht). RF: receptive field, VISp: primary visual cortex, FOV: field of view.

Selectivity of spiking responses for visual stimulus features such as spatially localized, moving, oriented edges is a hallmark of primary visual cortex neurons. We thus measured neuronal direction and orientation tuning curves using drifting gratings, as well as receptive fields using sparse noise stimuli (Fig. 4B, fig. S8A-B). Direction and orientation tuning involve intracortical interactions, while topographically organized (i.e. retinotopic) receptive fields reflect the fidelity of retino-thalamo-cortical mapping of visual space (*27*).

Ht neurons responded to drifting gratings with a broad range of individual direction and orientation preferences, and the same "salt and pepper" spatial organization as that of overlying cortical neurons (Fig. 4C). Regarding receptive field properties, Ht neurons were retinotopically organized, suggesting topographical retina-to-Ht mapping (Fig. 4D left). Moreover, within single anatomically-defined radial "pseudo-columns", cortical and Ht neurons displayed similar retinotopic preferences (Fig. 4D right, 4E), further suggesting that cortical and Ht neurons form integrated functional units. Hence, Ht neurons are essentially indistinguishable from primary visual cortical neurons in terms of their core responses to visual stimuli.

### Cortical substitution

Given the extent of the spatial and circuit organization of Ht neurons and the connections they form with the brain and spinal cord, we next examined whether Ht neurons could functionally substitute for cortical neurons in a behaviorally relevant context. To test this possibility, we designed a whisker stimulation task that requires cortical function and assessed whether Ht neurons could compensate for a decrease in perceptual performance upon optogenetic inhibition of cortical activity.

To be able to transiently inhibit cortical function, we first injected a soma-targeted inhibitory opsin-expressing AAV (pAAV-hSynI-STO-stGtACR2-FusionRed, (*28*)) in SSp-bfd (we used intrinsic optical signal imaging upon single-whisker stimulation of the B2 whisker to target the injection site, see Methods, Fig. 5A). We confirmed that blue light inhibited spiking in Cx stGtACR2-expressing neurons in acute brain slices (Fig. 5B). We then trained mice to signal the detection of whisker deflection by licking a water spout. Under baseline conditions, cKO mice had the same detection threshold as control mice, indicating similar detection skills (Fig. 5C). Deflection amplitude influenced responses: strong whisker deflections were more likely to be detected (i.e. resulted in a higher probability of licking) than weak ones (fig. 5D left). Varying deflection amplitudes allowed us to generate a psychometric curve that mapped the mice’s perceptual capacity. By interleaving trials of cortical inhibition using blue light with control sessions trials where light was projected outside of the cranial window, we first demonstrated the role of the cortex in whisker deflection perception: inhibition of cortical neurons impaired the ability to detect whisker stimulations in Ctl mice (Fig. 5D left, fig. S9A-B). In Ht mice, however, inhibition of cortical neurons had no effect on performance, (Fig. 5D right, fig. S9C). This result demonstrates that Ht neurons can functionally substitute for cortical neurons when silenced.

**Figure 5.**
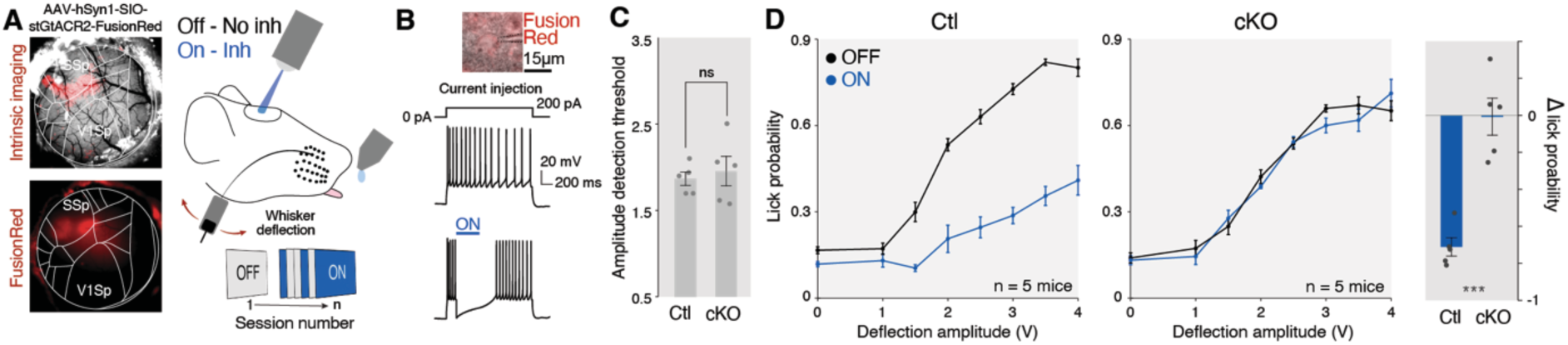
Ht neurons can functionally substitute for cortical neurons. **(A)** Left: Intrinsic optical signal imaging of SSp-bf upon B2 whisker stimulation to define the viral vector injection site (top). FusionRed immunofluorescence imaged through a cranial window reflecting neuronal populations expressing stGtCR2-FusionRed injection (bottom). Right: Schematic representation of the whisker deflection detection task. Optogenetic stimulation trials (during which the B2 barrel is inhibited) are interleaved with trials without light stimulation. **(B)** Optogenetic inhibition of stGtACR2-expressing neurons in acute brain slices. **(C)** Detection threshold in OFF conditions of Ctl and cKO mice (unpaired t-test, *P* = 0.76). **(D)** Licking probability of Ctl (left) and cKO mice under normal conditions (OFF - black) and blue light stimulation (ON - blue) for different whisker deflection amplitudes. Average relative lick probability loss, per mouse (unpaired t-test, *P* = 0.0002). Inh: inhibition.

Our results reveal that the cellular diversity and molecular identity, input-output connectivity, electrophysiology and function of the neocortex can be reproduced in a modified spatial context. Hence, there are multiple anatomical solutions to a specific behavioral task.

Structure duplication followed by divergence is a driving force in brain evolution (*5*, *6*), and is thought to have underlaid the emergence of basal ganglia and cerebellar circuits (*29*, *30*), bird and human vocal pathways (*6*, *31*), and higher-order cortical areas from primary sensory and motor ones (*32*, *33*). The specific design of a given species’ brain reflects an evolutionary-tailored, fleetingly optimal solution to anatomical developmental and functional constraints. Our findings provide an example of how loss of a single gene can lead to the duplication of a functional brain structure, with a distinct cytoarchitecture (i.e. concentric vs. laminar), but surprisingly conserved topography and connectivity. Although Ht circuits appear largely duplicated and redundant with original cortical circuits, with which they function in parallel, structural divergence with emergence of a new function could, in principle, develop along generations through select evolutionary pressure (*34*).

Subcortical Ht in humans, when clinically manifest, cause seizures and intellectual deficit, likely reflecting abnormal connectivity within the Ht and with adjacent structures (*35*– *37*). Ht can, however, present some level of functional activity in humans, as instances of co-activation of Ht with the overlying cortex upon motor or visual stimulation have been reported, suggesting that some level of self-organization also occurs in human Ht (*38–43*).

Ht neurons have preserved neuronal polarity despite grossly abnormal orientation, which is also occasionally observed in the cortex (*44*). Despite their abnormal location and orientation, however, the circuit connectivity of Ht neurons essentially duplicates that of cortical neurons. Our findings do not distinguish whether such duplication reflects cell-autonomous or non-cell-autonomous processes, since inputs to both structures are remarkably similar. However, they suggest that when attempting to reverse engineer neuronal circuits *in vitro*, as in organoids, priority can be put on understanding and emulating the molecular characteristics of the neurons comprising the circuits, rather than on recreating the exact spatial positioning of the neurons within these circuits. Altogether, these findings highlight a remarkable level of self-organization of neuronal systems as they emerge within host circuits: their ability to adapt to changing conditions while maintaining robust functional integrity.

## Acknowledgments

We thank the Genomics, the Human Cellular Neuroscience, the Bioimaging and the FACS platforms of the University of Geneva; the Advanced Lightsheet Tmaging Center of the Wyss Center Geneva; the CNRS-TAAM platform at Orleans, France; N. Baumann and J. Prados for assistance with bioinformatics analyses and for providing the DNN structure; L. Frangeul for assistance with whisker trimming experiments; all members of the Jabaudon laboratory for their comments on the manuscript and for constructive exchanges during the project, as well as inter-lab discussions; the PE and A2 INMED animal facilities, F. Michel from the INMAGTC imaging facility, and F. Bader and the rest of the PBMC molecular and cellular biology facility of INMED, Marseille, for technical support.

## Authors contribution

SRP and DJ conceived the project and designed the experiments, with the exception of those related to the morpho-functional organization of the barrel cortex, which were designed by FW, MM and JBM. SRP, MG, DS, AF, FW, SF, GM, EM, CGF, DF and EK performed the experiments, SRP performed the snRNA-seq and MERFTSH bioinformatic analyses. SRP, AF, FW and MG performed the *in* vivo tracing experiments. DS performed the *in vivo* recordings and analysis from the somatosensory cortex, and AF performed post-hoc histological confirmation of probe placement. MG, GM and CGF performed the *in vivo* experiments and analyses for the visual system and the optogenetic manipulation. SF performed the electrophysiological recordings on sections. FF generated and provided the mice line used in this study and their initial characterization. DJ and SRP wrote the manuscript. MG, DS, AF, SF, GM, EM, CGF, DF, EK, FF, MM, JBM, SEB and FW revised and edited the manuscript.

## Data and code availability

Analyses in this study were performed using existing code. The rest of the data supporting the findings of this study are available from the corresponding author upon reasonable request.

## Writing process using generative A1 and AI-assisted technologies

we used Chat-GPT to improve specific sections of the manuscript. After using it, we reviewed and edited the content as needed and bear full responsibility for the content.

## Materials and Methods

### Mice

All the experimental procedures described here were conducted in accordance with the Swiss laws and previously approved by the Geneva Cantonal Veterinary Authority. Swiss mouse husbandry was done at Charles River Laboratory and at the institutional animal facilities under standard 12h:12h light:dark cycles with food and water *ad libitum*. Mouse husbandry in France was carried out conforming to national and international directives (directive CE 2010/63 / EU, French national APAFIS n° 23424) with protocols followed and approved by the local ethical committee (Charles Darwin, Paris, France), the Institutional Animal Care and Use Committee of Aix-Marseille University [2020080610441911_v2(APAFIS n° 26835)] and following the guidelines of the French National Institute of Health and Medical Research (INSERM, provisional approval N007.08.01). Embryonic day (E) 0.5 (overnight mated females) was established at time of detection of vaginal plug. *Emxl:Cre* (*45*) and *Emlllox* (*9*) lines (C57BL6/J genetic background) were combined to generate a conditional deletion of *Emll*(cKO) in the dorsal pallium (*9*, *lO*). In this study we indistinctively analyzed males and females.

### Genotyping of transgenic mice

Biopsies were collected and DNA was extracted and amplified using the MyTaq Extract-PCR kit (meridian bioscience) following manufacturer instructions. The following primers were used for genotyping:

*Emx1:Cre* forward (CAACGGGGAGGACATTGA)
*Emx1:Cre* reverse (TCGATAAGCCAGGGGTTC)
*Control* reverse (CAAAGACAGAGACATGGAGAGC)
*Eml1lox* forward (GAAAACGTGCTTTGCTGTGTACATAGG)
*Eml1lox* reverse (CACCCACTGAAGAAATGACTGGCAG)

### Tissue microdissection, single-nuclei sorting and sequencing

Acute coronal brain sections (300 µm) were performed on a vibrating microtome (Leica, VT1000S) and SSp was microdissected with micro-scalpel using a Leica Dissecting Microscope (Leica, M165FC) in ice-cold oxygenated ACSF under RNase-free conditions. For cKO animals, we microdissected the somatosensory cortex (SSp) and the Ht region underneath. Brains from individual adults were microdissected separately, on ice. We pooled the microdissected regions of 2 Ctl animals and 3 cKO animals. Nuclei were isolated using the EZ Nuclei isolation kit. All the procedure was performed on ice and in RNAse free low-binding tubes. Briefly, tissue was resuspended in 2mL of ice-cold isolation buffer (Nuclei EZ prep, Sigma NUC101), dounce homogenized (KIMBLE Dounce tissue grinder, Sigma D8938) and incubated 5 minutes on ice in a total 4 mL volume. The dissociated nuclei were centrifuged (500 g at 4°C, 5 minutes), resuspended and incubated in EZ buffer for other 5 min on ice. We performed two washes with 1% BSA (bovine albumin serum) PBS (phosphate-buffered saline) containing 50 U/ml of SUPERase-In (Thermo Fisher, cat# AM2696) and 50 U/ml of RNasin (Promega, cat# N2611). Finally, nuclei were filtered through a 30 µm strainer and stained with Hoechst (Invitrogen H3570, 1:500) for 5 minutes. In order to remove debris and only retain good quality nuclei, we used fluorescence-activated nuclei sorting (FANS) on a Beckman Coulter MoFlo Astrios. Approximatively 20k nuclei were sorted and 42 µl of nuclei suspension was used to load the 10x Genomics snRNA-seq preparation, according to the manufacturer’s instructions (10X Genomics Chromium 3’ Gene Expression Kit). The cDNA libraries were quality controlled using a 2100 Bioanalyzer and TapeStation from Agilent, and sequenced using a HiSeq 2500 sequencer from the iGE3 platform at the University of Geneva. FASTQ files from sequencing were used as inputs to the 10X Genomics Cell Ranger pipeline (v3.0.2) with pre mRNA transcript detection of GRCm38 mouse genome.

### Tissue collection and MERFTSH tissue processing

MERFISH procedure was performed following instructions from VIZGEN MERSCOPE Platform. A cKO brain was collected in cold ACSF. The obtained tissue was rapidly fresh-frozen in cold isopentane and preserved at -80°C. One single 10 µm coronal section, using a Leica CM3050 cryostat, was collected on a MERSCOPE slide. Briefly, sections were postfixed for 10 minutes in fresh 4% PFA and preserved for 24 h in 70% EtOH. This was followed by the 463 gene panel (see table S1) incubation overnight at 37°C. After hybridization, the section was embedded and cleared. Finally, once the specimen in the MERSCOPE instrument from the Human Cellular Neuroscience Platform at the Geneva Campus Biotech, we ran the acquisition pipeline (https://vizgen.com/).

### Analysis

All single-cell transcriptomics analyses were performed using R Statistical Software (v4.0.2), the Seurat package (4.0.1) and Bioconductor packages. Graphs and visualization were generated with ggplot2 (3.4.4) and ComplexHeatmap (2.6.2).

#### Single-nuclei RNA sequencing

The three generated libraries (Ctl, Cx and Ht, see Fig. 1 and fig. S2) were merged as a single Seurat object and processed for quality controls following the same standards. Only nuclei with more than 1000 detected genes, more than 2’500 detected RNA molecules and less than 1% of mitochondrial RNA expressed genes were retained. In order to detect and exclude possible doublets, we ran the DoubletFinder package (2.0.3). We then performed: (1) gene count normalization to the total expression and log-transformation, (2) highly variable genes detection and principal component analysis, (3) graph-based clustering (30 first PCAs and a clustering resolution of 0.5), (4) UMAP calculation.

#### DNN cell-type prediction

Cell identities were predicted using a feedforward neural network trained on an adult single-cell transcriptomics dataset (*46*). Cells from the SSp, sequenced with the 10x Genomics Technology, were split into training (40’425 cells), validation (20’210 cells) and test (20’407 cells) datasets. The model was trained to classify cells into "subclasses" as annotated in the original dataset. These subclasses are composed of 10 glutamatergic categories (L2/3 IT, L4 IT, L4/5 IT, L5 IT, L5 PT, L6 IT, L6 CT, L6b, L5/6 NP and Car3), 6 GABAergic (Vip, Lamp5, Pvalb, Sncg, Sst and Sst Chodl) and 5 non-neuronal (Astro, Oligo, Endo, Micro-PVM, SMC-Peri). The prediction of the model on the test dataset, composed of cells unseen by the model, yielded good prediction performances: 97.8% of the test cells were correctly predicted. In more detail: L2/3 IT: 99.7% - L4 IT: 98.2% - L4/5 IT: 96.1% - L5 IT: 90.1% - L5 PT: 100% - L6 IT: 96.3% - L6 CT: 98.7% - L6b: 99.4% - L5/6 NP: 99.5% - Car3: 100% - Vip:99% - Lamp5: 97.9% -Pvalb:98.1% - Sncg: 98.3% - Sst: 99.6% - Sst Chodl: 100% - Astro: 100% - Oligo: 100% - Endo: 100% - Micro-PVM: 100% - SMC-Peri: 100%. All the cells of the current study were predicted simultaneously. The three samples displayed high prediction rates and we did not observe sample bias (fig. S2C). DNN prediction confidence scores were calculated as the difference between the best cell-type prediction value and second-best cell-type prediction value for each cell.

#### Molecular distance

To calculate the molecular distance between cell-types across the first 30PCAs, we performed the following calculations: (1) established the mean position of single cell-types in each PCA space for Ctl, Cx and Ht samples. (2) We measured the euclidean distance in each PCA space between conditions (dist() function from the stats package) and (3) we computed the mean of the calculated distances in order to obtain a single distance value represented as a heatmap (ComplexHeatmap package 2.6.2).

#### MERFISH data processing

Gene expression and coordinates matrices were obtained from the instrument output. They were assembled as a Seurat object and briefly, we kept only cells with more than 15 detected genes, we normalized the expression to the total expression and performed a log-transformation, detected highly variable genes, performed principal component analysis, graph-based clustering (30 first PCAs and a clustering resolution of 0.5) and finally UMAP calculation. Cell types were annotated manually based on the expression of well-known cell-type markers included in the used gene panel. Spatial subsets (Cx and Ht) were manually done using the Gatepoints package (0.1.4).

#### Medio-lateral modelling

We subset the Cx and the Ht separately and defined the glutamatergic neuronal position using the *principal_curve* function of the devtools package (2.4.1) as depicted in fig. S5A. Cx neuronal position score was binned in 7 segments (750 cells per bin), used to reconstruct medio-lateral positioning using an ordinal regression model with the bmrm package (4.1) as previously described (*47*). The regularized ordinal regressions were used to separate medio-lateral segments (1 to 7). During modelling, each gene is assigned a weight based on its relevance for correct cell position prediction. The 35 most negatively and the 35 most positively weighted genes were retrieved as core genes and used for Ht cells medio-lateral score prediction (see table S2).

#### Transcriptional waves

Glutamatergic neurons were aligned based on their medio-lateral position and their expression of the medio-lateral model genes was corrected using weighted expression in neighbor cells. This was performed separately for Cx and Ht. All transcriptional waves were normalized to their maximum value.

### Immunofluorescence

Mice were transcardiacally perfused with 4% PFA/Phosphate-buffer (PB 0.2M) under thiopental anesthesia, brains extracted and post-fixed overnight in 4%PFA/PB at 4°C. Brains were subsequently sectioned at 80 µm using a vibrating microtome (Leica, VT1000S).

#### lmmunofluorescence on free-floating sections

sections were incubated 30 minutes in 1x antigen retrieval solution (Sigma-Aldrich C9999) at 80°C and washed three times in 1x PBS. They were then incubated 30 minutes in blocking solution (3%BSA, 1x PBS and 0.5% Triton X-100) at room temperature. Sections were incubated with primary antibodies (see list below) over two nights at 4°C. After three 1x PBS washes, sections were incubated with respective secondary antibodies (1:500) and Hoechst (1:2000) during 2 hours at room temperature.

#### Flatten brain preparations

P10 mice were transcardiacally perfused with 4% PFA/Phosphate-buffer (PB 0.2M), under tiletamine/zolazepam (Zoletil, 40 mg/kg) and medetomidine (Domitor, 0.6 mg/kg) anesthesia. Brains were immediately extracted and cortices dissected, flattened between microscope slides and post-fixed in 4% PFA overnight. Cortices were included in 4% agar and tangential sections were cut at 100 µm on a vibrating microtome (Leica, VT1000S).

#### Antibodies

rabbit anti-CUX1 (1:400, sc-13024), rat anti-BCL11B (1:500, ab18465), mouse anti-RORB (1:500, PP-N7927-00), rabbit anti-AnkG (1:1000, sc-28561), guinea-pig anti-VGLUT2 (1:2000, ab2251), rat anti-SST (1:200, AF2727), goat anti-PROX1 (1:400, MAB354), mouse anti-REELIN (1:1000, ab78540), mouse anti-PVALB (1:500, PV235). Adequate secondary antibodies were all used at 1:500 (Life Technologies).

#### EdU pulse and revelation

250µL (100µL/10g of animal) of EdU (7.5mg/mL) were injected intraperitoneally in pregnant damns at the indicated embryonic days. For its revelation, we mixed in the following order 1x PBS (429 µL) with 20µL of 100mM CuSO_4_, 1µL of AF azide 647 and 50µL of 1M Sodium L-Ascorbate. The mix was immediately added to free-floating sections (80 µm) and incubated at room temperature for 30 minutes. This procedure was performed after the immunofluorescence protocol described above.

### Cranial window and microprism implantation

Pain management was performed by first administrating Buprenorphine subcutaneously (0.1 mg/kg) before starting the surgery and a local anesthetic right before the surgery (mix Lidocaine/Bupivacaine, 6 mg/kg and 2.5 mg/kg respectively). Mice were anesthetized with isoflurane and then placed either in a stereotaxic apparatus (Stoelting Co.) and a Nanoject II (Drummond, #3-000-026) or a stereotaxic apparatus (Model 940, Kopf) with a custom-made nose-clamp, adapted to maintain the position of the animal and allowing head rotation. Body temperature was constantly monitored through a thermic probe and adjusted to -37°C via a heating pad placed below the mouse (DC Temperature Controller, FHC). At the end of the surgery, an anti-inflammatory was administered subcutaneously (Carprofen, 7.5 mg/kg) and animals were warmed with a heating lamp for at least 15 minutes until recovery from anesthesia. A second anti-inflammatory (Ibuprofen, Algifor) was added to drinking water for 3-days post-op and the weight was checked daily to ensure that weight was kept above 15% of the original weight prior to the surgery.

#### Cranial implant

Animals were first implanted with a cranial implant. After removing the skin and tissues on top of the head, the skull was cleaned, dried and thinned. A custom-made metallic implant was placed on the top of the skull using a custom-made holder. It was then fixed with a layer of glue (Loctite 401, Henkel) and additional layers of dental acrylic (Pala, Kulzer) to solidify the implant. Dental acrylic was covered with black nail polish to prevent light contamination from visual stimuli during experimentation.

Cranial window surgery: Animals were implanted with a custom-made glass window composed of a bigger top round cover slip of 7 mm diameter respectively and two superimposed 5 mm diameter cover slips, respectively. The three concentric cover slips were glued together with UV glue (Optical Adhesive n°68, Norland). The craniotomy was the size of the smaller window and was drilled around the region of interest. Before removing skullcap, we used a custom-made perfusion chamber with saline solution (NaCl 0.9%) to rinse the craniotomy continuously and reduce bleeding stains. A custom-made holder with air-suction was used to hold and position the window. The cranial window was gently brought down until it was in contact with the brain. The window was then fixed with UV glue, super glue and dental acrylic.

#### Microprism implantation surgery

For microprism implantation, a glass prism with aluminum coating (1mm side, Tower Optical Corporation) was UV glued below the cranial window. Following craniotomy, the dura was entirely removed from accessible brain surface. A small incision was performed with a blade attached to the holder at the exact prism insertion site. The microprism assembly (window and prism) was then inserted inside the cortex and the window was attached as described in the previous section. Mice could be imaged after at least two weeks of recovery from surgery (*48*).

### Stereotaxic injections

Coordinates were calculated from bregma and defined for each experiment. The respective coordinates will be detailed below. Surgeries were performed as described above. Glass pipettes (5-000-2005, Drummond) were pulled (Model P-97, Sutter Instrument Co) and further broken to obtain a tip of -10-15 µm inner diameter and further beveled to create a sharp tip and avoid cortical damage during insertion.*Callosal projections tracings*: The skull was exposed by an L-shaped section on the head skin to expose the bregma and opened with a drill (outer diameter -60 µm). Cholera toxin subunit B coupled to Alexa 555 (CTB555, ThermoFisher, c34776) was injected medially was injected medially (Ctl - AP +1.0, ML +0.5, DV -0.6 - cKO - AP +1.0, ML +0.5, DV -0.8) and CTB coupled to Alexa 647 (CTB 647, ThermoFisher c34778) was injected laterally (Ctl - AP +1.0, ML +1.2, DV -0.6 - cKO – AP +1.0, ML +1.2, DV -0.8). The injection volume of each CTB was approximatively of a 100nL. The skin was sutured and mice quickly recovered after surgery on a heating pad. Brains were processed 7 days after the injection date.

#### MOp and SSs projections tracings

again, these tracings were performed combining CTB555 and CTB647. The same procedure that for CC tracings was followed. Instead, CTB 555 was injected in MOp (Ctl - AP +1.34, ML +1.8, DV -0.6 - cKO - AP +1.34, ML +1.8, DV -0.75) and CTB 647 was injected in SSs (Ctl - AP -0.9, ML +5.0, DV -0.6 - cKO - AP -0.9, ML +5.0, DV -1). Brains were processed 7 days after the injection date.

#### Spinal cord retrograde injection

in adult animals, the cervical spinal cord was surgically exposed and 220nl of rAAV2-retro-CAG-TdTomato (vector core facility of the University of North Carolina) were injected in both sides of the cervical corticospinal tract (segment C2/C3). The skin was sutured and mice quickly recovered after surgery on a heating pad. Tracings were performed in 3 Ctl and 3 cKO mice. Brains were processed 10 days after the injection date.

#### Ht anterograde injection

Similarly, we exposed the skull by a longitudinal section of the head skin to expose the bregma. 200nL of an antero rAAV2-CAG-Gfp (vector core facility of the University of North Carolina) were injected in the Ht (Bregma: AP -0.9, ML +1.0, DV -0.8). After skin suture, mice were put on a heating pad until recovery. Tracings were performed in 2 cKO mice. Brains were processed 10 days after the injection.

#### Thalamus anterograde injection

P0-P1 mice were anesthetized by hypothermia on ice for 8-10 min and then placed on a pup stereotaxic adaptor (Stoelting, 51615), pre-cooled with dry ice, and fixed to a stereotaxic frame associated to a digital console (Kopf instruments). Coordinates were established using the Lambda point (crossing of transverse and superior sagittal sinus) as a reference. The skin was aseptized with Vetadine and 70% ethanol. A small incision was made in the scalp followed by a gentle drill at the exposed skull location. 2.3 nL of AAV1-CAG-tdTomato (59462-AAV1; Addgene) were injected twice, 1 minute interval, in the VPM (AP +0.4 mm, ML +1.0 mm, DV +2.3 mm). Pups were then placed on a heating pad to restore a proper body temperature and given back to the mother. Tracings were performed on 1 Ctl and 4 cKO mice. Brains were processed 7 days after the injection.

#### Intrinsic optical imaging (IOS) guiding

To assess the injection sites of experiments described in Fig. 4 and Fig. 5, we used a custom-made wide-field epifluorescence microscope setup including a sCMOS camera (ORCA-Flash4.0 V3, Hammamatsu). Magnification was determined through a 0.63X C-mount camera adapter for Olympus Microscopes. The field of view size was 5.6 mm x 5.6 mm. The camera and adapters were mounted on a base allowing vertical movement with manual focus. LED white illumination (740mW, 957 1225mA, Thorlabs) could be controlled via a T-Cube LED driver (LEDD1B, Thorlabs). To record intrinsic signals, Cy3/5 excitation, emission and dichroic filters were used. Objective (MVX Plan Apochromat with 2x, Olympus) was attached to the microscope base. The somatotopic mapping consisted in repetitive rostro-caudal pulsatile deflections (-1 963 mm amplitude) of either B2 or C2 whiskers for 50-90 trials followed by quiet window. The retinotopic mapping protocol consisted of drifting bars. A contrast reversing checkerboard was presented within the bar to better drive neural activity. In each trial the bar was swept in the four cardinal directions: left to right, right to left, bottom to top, and top to bottom. Single trials were repeated in average 15-30 times (*49*). Mice previously implanted with a 5 mm diameter cranial glass window were head-fixed on the platform and anesthetized during the procedure with isoflurane level lowered to 1%. Body temperature was monitored with a probe and adjusted to 37°C using a heating pad (DC Temperature Controller, FHC).

#### VlSp GCaMPf6 injection

we performed several injections of an AAV-syn-GCaMP6f virus (-100 nL each) in the middle of VISp, with different depths (800, 500 and 200 µm) to target both Ht and Cx neurons.

#### SSp opto injection

we performed two injections of an AAV-hSyn-stGtACR2-FusionRed virus in the center of the B2 whisker signal in wS1 and wS2 respectively. A single depth (100 µm) was targeted to get only the Cx neurons infected. Injections were done using a single-axis oil hydraulic micromanipulator (R.MO-10, Narishige). The pipette was slowly inserted inside the cortex until reaching the desired depth. Viruses were injected at a speed of -2nL/s. When the whole volume was injected, we waited 5 minutes with the pipette in the same position before gently removing it.

### Tissue clearing and whole brain imaging

Mice were transcardiacally perfused with 4% PFA/PBS and brains were postfixed with 4% PFA/PBS at 4°C overnight. Subsequently, brains were clarified following the CLARITY protocol (*50*, *51*). Imaging of entire clarified brains were acquired using a light-sheet microscope (MesoSPIM (*52*)) with Z stacks at 5 µm spacing and a zoom set at 0.8x resulting in an in-plane spatial resolution of 8 µm (2048×2048 pixels, further specifications under https://www.campusbiotech.ch/en/facilities/imaging-and-microscopy). Tissue clearing and 3D imaging for retrograde tracing experiments was performed for AAV Ht anterograde and corticospinal retrograde labelling. For fig. S1A and Fig. 2C images, we isolated the heterotopia and the CST respectively by manual segmentation using Imaris (v9.9, BITPLANE). Using the same software, for Fig. 2D and fig. S6A, we performed digital sagittal and coronal sections of the imaged brains using the clipping plane tool.

### Electrophysiology on acute brain sections

#### Electrophysiological characterization

Three-hundred-micrometer-thick acute coronal slices were prepared from adult Ctl and cKO mice, previously injected, as aforementioned, with an rAAV2-CAG-TdTomato in the spinal cord 10 days before. Slices were kept for at least 30 min in modified artificial cerebrospinal fluid (aCSF) at 33°C (87mM NaCl, 2.5 mM KCl, 7mM MgCl_2_, 0.5mM CaCl_2_, 1.25mM Na_2_HPO_4_, 25mM NaHCO_3_ and 5 mM glucose), supplemented with 1mM kynurenic acid (Sigma K3375) and oxygenated with 95% O2 and 5% CO2 which was gradually replaced by the the aCSF used for patch-clamp recording (125 mM NaCl, 2.5 mM KCl, 1mM MgCl_2_, 2.5mM CaCl_2_, 1.25 mM Na_2_HPO_4_, 26 mM NaHCO_3_ and 11 mM glucose, oxygenated with 95% O2 and 5% CO2). The slices were then transferred in the recording chamber, submerged, and continuously perfused with aCSF. Patch-clamp recordings were performed on Td-Tomato^+^ retrogradely labeled L5B neurons in the primary motor cortex area. The internal solution used for the experiments contained 140mM potassium methansulfonate, 2 mM MgCl_2_, 4 mM NaCl, 0.2 mM EGTA, 10 mM HEPES, 3mM Na_2_ATP, 0.33mM GTP and 5mM creatine phosphate, 0.3% biocytin; pH 7.2; 295 mOsm. In voltage-clamp configuration, the voltage was clamped at -60mV. Access resistance was monitored by a hyperpolarizing step of 14 mV. Ih current was calculated following a hyperpolarization step of -40mV for 500ms. The neuron was then placed in current clamp mode and the resting membrane potential was monitored every 10 s and averaged for 5 consecutive acquisitions. Four steps of 50, 100, 200 and 400 pA were applied for 500 ms. During the recording, neurons were passively filled with biocytin and at the end of the recording, the patch pipette was slowly retracted from the cell membrane to obtain an outside-out patch in order to maintain the integrity of the plasma membrane for further neuron morphology analysis. Fifteen minutes after the end of the recording, slices were fixed with 4% paraformaldehyde at 4°C overnight and incubated with Alexa 647 coupled-streptavidin (Invitrogen S21374, 1:500 in PBS-10% tween) for 6 h at room temperature. Sections were then washed with PBS before mounting. All analyzed neurons were recorded from at least 3 different animals per condition.

#### Inhibitory opsin experiments

To validate the behavioral use of AAV-hSyn1-SIO-stGtACR2-FusionRed, we performed acute slices from 2 adult cKO mice injected with the virus in the SSp-bf 30 days before. Patch-clamp recordings were performed on FusionRed^+^ neurons. In current clamp configuration, a depolarizing step of 200pA was applied for 1450 or 4500 ms. 200ms following the beginning of the current injection, a continuous blue-light pulse was applied for 400ms or 3s respectively through the 60X objective. Liquid junction potential was not corrected. Recordings were amplified (Multiclamp700, Axon Instruments), filtered at 5 kHz and digitalized at 20 kHz (National Instrument Board PCI-MIO-16E4, IGOR WaveMetrics), and stored on a personal computer for further analyses (IGOR PRO WaveMetrics). All analyzed neurons were recorded from 2 independent cKO mice.

### Single whisker stimulation and c-Fos immunostaining

Adult isoflurane anesthetized mice underwent complete whisker trimming of the right whisker pad and of all whiskers but one (C2) on the left side. After anesthesia recovery on a heating pad, mice were placed in an enriched cage for 1.5h. Immediately after, mice were transcardially perfused with 4% PFA/PBS and brains were postfixed with 4% PFA/PBS at 4°C overnight. As previously described, brains were sliced and brain sections were treated for antigen retrieval, incubated with a mix of rabbit anti-c-Fos (1:5000, sc-52) and mouse anti-RORB (1:500, PP-N7927-00) overnight at 4°C and with the due secondary antibodies. We performed this experiment on 3 Ctl and 3 cKO mice.

### Electrophysiology in vivo

#### Intrinsic optical imaging (IOS) guiding

used to locate a specified barrel in somatosensory cortex in vivo. IOS was recorded using a video acquisition system. The camera was positioned orthogonal to the exposed surface of somatosensory cortex. Video images of the illuminated cortex were acquired at 50 frames per second with a CCD camera (QImaging). An image of the superficial vessel topography was made using high contrast green light (528 nm). For IOS, under infrared illumination (860 nm) the camera was focused 100-400 µm below the cortical surface to minimize surface artifacts. The specified whisker was stimulated at 5 Hz for 10s following a 5s prestimulus period (used as background). Trials were repeated 10-20 times with 1 minute between trials. For analysis, the average pre-stimulation image was subtracted from each subsequent recorded image within a block of trials. Change in the scattering was accumulated through all trials. To target the principal barrel for recording in the somatosensory cortex, IOS response was superimposed on the vessel map.

#### Extracellular recordings

performed in P8-45 rats under urethane (1 g/kg) anesthesia following preparations described previously (*53*). The electrodes were placed in the centre of the barrel visualized using IOS as described above. Multisite silicon probes (Neuronexus, USA) of 16 channels with a 100 µm separation distance were inserted to the depth of 1.5 mm for simultaneous recordings of activity both in heterotopia and barrel cortex. The single whiskers were stimulated by piezo actuators (Noliac).

#### Stimulation

before the experiment, whiskers were trimmed to a length of 4 mm. A loop from the wire of 0.2 µm diameter was glued to the end of piezo benders (Noliac), and the tip of the whisker was inserted 2 mm into the loop, so that the whisker rested snugly inside. To induce deflection of the piezo benders, square 40-70 V pulses of 10 ms duration were applied. To avoid depression of the evoked response, whiskers were alternately stimulated at 30 s intervals. Cortical responses to 50-100 stimuli were recorded for each whisker. To characterize cortical receptive fields the principal whisker (PW) and first-order adjacent whiskers (AW) were also stimulated, (first-order adjacent whiskers being whiskers neighboring the principal one).

#### Analysis

to characterize the evoked neuronal activity in the barrel cortex and heterotopia, the multi-unit activity (MUA) was used. Using custom-written functions in MATLAB 2016a (MathWorks), raw data were explored to detect MUA. The analysis was conducted in a 15-s windows after the stimulus to define evoked activity. MUA was detected in a band-passed signal (>200 and <4000 Hz) in which all negative events exceeding 3.5 SDs calculated over the entire trace were considered as spikes (>99.9% confidence;). Detection of the MUA onset time was done using the accumulation approach as described previously (*23*).

### Behavioral training and optogenetics

#### Behavioral training

A Matlab (MathWorks, USA) custom-made graphical user interface (GUI) was developed to control the behavioral tasks and allow for real time monitoring of animal performance. Animals underwent water restriction 2 to 4 days before training started and were handled every day for at least 10 minutes. During pre-training phase, mice were habituated to head-fixation and placed on the setup for about 15 min and then for longer durations. On the first sessions of training, mice were exposed to the task with trials where water was automatically delivered following Go trials (rewarding with 100% reward probability) to engage licking behavior. As soon as they started licking spontaneously, automatic water delivery was stopped and water was only delivered if mice licked for Go trials during the response window. This first phase of training typically lasted for one or two sessions. Go trials proportion was progressively reduced as Catch trials (non-rewarded) proportion was increased in the following sessions. Mice were trained once per day every day at the same hour until they displayed stable behavior measured by their lick probability for all conditions and discrimination performance. Body weight was monitored daily and was kept above 80% of the weight measured prior to water- restriction. Mice received at least 1 mL of water per day, either during behavioral training or in their home cage. After 20 days of water-restriction schedule, mice were given access to water ad libitum for at least 2 consecutive days.

Mice were trained to a single whisker detection behavioral task under a water-restriction schedule. All right whisker-pad whiskers except for the B2 were trimmed close to their base with surgical scissors under a magnification binocular microscope. This operation was performed under 2% isoflurane anaesthesia before behavioral training. Mice were head-fixed on a platform and the B2 whisker was inserted inside a glass capillary tube (Wiretrol® II 5 & 10 uL, Drummond Scientific Company) placed 2mm away from the mouse’s snout, attached to a piezo actuator (Bimorph bendor piezo actuator PB4NB2S, Thorlabs) creating a small deflection of the whisker along the rostro-caudal axis. All tactile stimuli were generated through Matlab data acquisition toolbox controlling a piezo controller amplifier (3-Channel, Open-Loop Piezo Controller, MDT693A, Thorlabs) through a National Instrument card (NI PCIe-6321 and BNC-2110 Connector Block, National Instrument). Whisker stimuli consisted of one sine waveform pulse lasting 40 ms. The amplitude of the stimuli was varied across 7 amplitude values (1, 1.5, 2, 2.5, 3, 3.5, 4V). resulting in a tube displacement which varied between 200um and 800um (a 0.5V amplitude corresponding to a 200 µm capillary tip deflection). This allowed us to obtain psychometric curves of the detection performance as measured by their lick probability for each individual amplitude value.

The total trial duration of a single trial was 9 s: each trial started with a 2 s quiet window of 2 sec during which any detected lick resulted in trial abortion. This helped to reduce spontaneous licking. Following the quiet window, tactile stimulation was presented. From stimulus onset, the mouse was allowed to lick during a 2 s response window to indicate detection. After the response window, a consumption window of 5 s allowed the mouse to collect the reward. Each trial was followed by an inter-trial interval of at least 4.5 s. During a Go trial, licking the spout upon B2 whisker stimulation was associated to a drop of water reward of approximately 10 µL: "Hit trial". Failure to lick would result in a "Miss trial". In some trials, tactile stimulus presentation was omitted: "Catch trials". During Catch trials, the mouse had to refrain from licking (Correct rejection trials, CR) but if the mouse licked during catch trials it was punished with a time-out of 12 s (False Alarm trials, FA). If the mouse was not following the temporal structure of the task by licking during the quiet window too frequently, a 12 s early lick time-out punishment could also be applied, delaying next trial initiation. The proportion of each trial type was the following: Go trials = 70%, Catch trials = 30%.

#### Atlas fitting and registration

The reference Allen Institute mouse brain atlas projected to match the skull tilt in our experiments and was manually fitted to the wide-field and DLP projector field-of-view of each mouse. This was done by visually aligning the atlas boundary lines to reproducible landmarks in the somatotopic and retinotopic maps obtained in wide-field. Landmarks used include the outlines of the sign map patches (marking distinct visual areas) and the position of B2 whisker activity bumps in S1 and S2.

Placement of the light pattern for optogenetic silencing experiment was done using information from wide-field functional mapping, fluorescence imaging and vessel patterns as anatomical landmarks. Final placement of the pattern was done using a Matlab custom-made graphical user interface (GUI) developed from the Matlab App Designer in order to realign single day anatomical image to reference image as to correct for potential shift in the animal’s position.

#### Optogenetic inactivation experiment

Optogenetic silencing experiment were performed during behavior task performance using Ctl (n=5) and cKO (n=5) head-fixed mice. Animals of both groups were implanted with head fixation headposts and 5 mm cranial windows and injected with AAV-st-GtACR2. The stereotaxic virus injections were targeted based on the information obtained from wide-field functional mapping to obtain local expression of FusRed-st-GtACR2 in the C2-encoding locations of S1 and S2 somatotopic maps. The blue light activated inhibitory opsin St-GtACR2 inducing shunt-inhibition by hyperpolarizing membrane potential was used to silence excitatory neurons in the cortex. Behavioral effect of the optogenetic inhibition of somatosensory neurons using the blue-light sensitive chloride channel stGtACR2 were also corroborated through in vitro electrophysiology experiments (Supp). In order to excite st-GtACR2 and obtain local wS1, wS2 inhibition during the experiment, a light pattern tailored to the injection sites was projected onto the cortical surface using a custom-made optical column with a DLP Projector system and 460nm high power LED (STAR 3.0 EVM Monochrome LED Projection Modules, DLP6500, ViALUX GmbH). The DLP projector system was used to deliver blue light at the surface of the cortex through the cranial window. It was controlled in real time by the Matlab custom-made graphical user interface to induce neuronal inhibition in a subset of trials (GUI). During the whisker detection task, blue light stimulation was delivered in 50% of trials interleaved with control trials. A double circular light pattern targeting the B2-encoding regions in wS1 and wS2 was used as optogenetic stimulation ("Opto" conditions). A simple circular pattern projected outside the cranial window was also included as a control condition ("NoOpto" conditions). Individual circular light pattern diameter at the cortical level was 600 µm. Within a session, 50% of the trials were pseudo-randomly chosen as "Opto" and 50% as "NoOpto" trials. Both patterns were projected during both Go trials and Catch trials. In all experiments, the intensity of the LED power was set between 0.05 W (2m W/mm^2^) and 0.145 W (7.5m W/mm^2^). The projector refresh rate was 100Hz and the duty cycle was 1sec allowing for continuous illumination. The duration of the projection was 4 s per trials. Projection started at trial initiation and lasted during the 2 s quiet window, the tactile stimulus presentation, and the 2 s response window. In case of trial abortion, illumination stopped immediately. To ensure that mice did not directly perceive the blue light used for optogenetic silencing, we isolated the optical column and the mouse skull with an opaque sleeve.

### Tmage acquisition, analysis and quantifications

All images were acquired at the Bioimaging platform of the University of Geneva on an upright Leica STELLARIS 5 microscope, using a HC PL FLUOTAR 10x/0.30 Dry or a HC PL APO CS2 20x/0.75 Dry objective or at the INMAGIC imaging facility with a Zeiss Z2 upright microscope equipped with a fully motorized fluorescence structured illumination module using either 5x, 10x or 20x dry objectives. All images were imported to Image J (2.9.0) using the Bio-Formats plugin. Biocytin filled neurons were imaged at the Human Cellular Neuroscience Platform of the Geneva Campus Biotech on a Nikon AxR (Galvano mode) with a 20x CFI Plan Apochromat Lambda D for morphology reconstruction and with an Eclipse 90i epifluorescence microscope (Nikon) to register their position on the section.

*Inhibitory neuron*s quantification: populations were quantified in the SSp of Ctl animals and cKO mutants. We counted the number of positive somas in a defined ROI and measured the density in the Ctl SSp, the Cx SSp and the putative Ht SSp. For each condition, we generated the mean density over 3 sections of a single mouse and at least 3 mice were quantified per condition.

#### Neuronal orientation

for this purpose, we used AnkG immunostainings, labelling the axon initial segment (AIS). Using Fiji (2.9.0), we measured the angle between the AIS and the white matter in the Cx and the Ht shell and the Ht core. We quantified 486 Cx, 594 Ht shell and 515 Ht core angles on 1 section from 2 different cKO animals. We then used ggplot2 to represent the AIS angles, for each of the analyzed conditions.

#### CC tracings

on images from cKO traced animals, we quantified the number of medially and laterally projecting retrogradely traced neurons over 3 different sections of each single animal (n = 4). Also, we measured their relative mediolateral position, defining the most medial part of the CC as 0 and the most lateral part of the heterotopia as 1. We then computed density plots for each of the quantified animals using ggplot2.

#### SSp projections retrotracing from MOp and SSs

we quantified the number of MOp, SSs, or dual projecting neurons in 3 different sections of each animal (n = 3 Ctl and n = 2 cKO). The positive cells were quantified in the Ctl SSp, in the Cx SSp and the Ht SSp. Proportions were quantified based on the total number of quantified cells per mouse.

#### Neuronal reconstruction and morphology analysis

Biocytin filled imaged neurons were imported in Imaris and neuronal morphology was manually reconstructed using the filament tool. Simultaneously, epifluorescence images were annotated in QuPath (*54*) for neuronal position and orientation. After, Imaris 3D confocal neuronal reconstruction files and QuPath annotated widefield images were further processed using a custom-made MATLAB R2022b (The MathWorks) graphical user interface. In brief, the maximum intensity projection of a confocal stack was aligned with its corresponding widefield image through manual control point pairing to infer an affine transformation. This alignment allowed the orientation of the reconstructed neuron along the XX and YY directions. Subsequently, the 3D neuronal arborization was resampled using evenly spaced vertices. Finally, the Cartesian coordinates of these vertices were transformed into spherical coordinates. Therefore, the presence of apical or basal dendrites at a given distance from the soma, as well as the tropism and flatness of these dendrites, were quantified respectively by considering the probability density functions of the radial, azimuthal, and elevation components. The three empirical measurements were followed by kernel density estimations.

#### PCA analysis on electrophysiological recordings

data for each recorded parameter were compiled to create a matrix of 54 neurons (21 Ctl, 20 Cx and 13 Ht) and 9 parameters: Vm, R-input, Ih, Rheobase, Burst-index, AP threshold, AP amplitude, AP halfwidth and fAHP. After normalization to the mean of each parameter and a scaling step, we performed a PCA analysis. To use the dataset from the rest of cortical glutamatergic types (*21*), we reduced the number of parameters to 5: Vm, R-input, Rheobase, AP threshold and AP amplitude. Also, as dataset is composed of mean values for single cortical-types, we computed the mean value for Ctl, Cx and Ht of each of the aforementioned parameters.

#### Two-photon imaging

To assess whether cortical visual responses are comparable for Cx and Ht neurons (Fig. 4), we imaged two cKO mice after they underwent stereotaxic surgeries and viral injection of GcAMP6f using a two-photon microscope. The two-photon microscope was custom-made (INSS Company). It consisted in a laser with wavelength range 690-1040nm (Tunable Ti:Sapphire with dispersion compensation MaiTai DeepSee, Spectra Physics) which beam was displaced with Resonant/Galvo scan mirrors (INSS) and the emitted signals were detected with 2 GaAsP amplified PMTs (PMT2101/M, Thorlabs). Additionally, a z-piezo (P-725 PIFOC Long-Travel Objective Scanner for fast z-scanning, Physik Instrumente) was used for volume imaging. Imaging was performed through Scanimage (Vidrio Technologies). Images were acquired at approximately 30 frames per second. The protocol used was the same as in description for wide-field imaging section. Mice were head-fixed in the two-photon setup with an LCD monitor (20×15cm, 60Hz refresh rate) positioned 10 cm from the right eye, with a 30° angle to the right of the midline. Visual stimuli were passively presented on that screen on the top of a gray background and resulted in a mix of sparse noise stimuli and drifting gratings stimuli. For the sparse noise stimuli, the screen was divided into a grid of 6*7 possible positions, respectively 6 positions along the vertical axis and 7 positions along the horizontal axis. Two polarities were tested (black and white), resulting in 6*7*2 possible stimuli. A square was 171*171 pixels, with 20.53 pixels per degree. The spatial frequency of a sparse noise stimulus was 0.06 cycles per degree. Each stimulus had a duration of 1 sec. For the drifting grating stimuli, 8 possible drifting gratings could be presented, with an orientation going from 0° to 360° with a 45° shift between each grating. Each stimulus had a duration of 2 sec. The spatial frequency of one drifting grating was 0.06 cycles per degree with a temporal frequency of 2 Hz. All single stimuli were presented exactly 20 times each. Sparse noise stimuli and drifting gratings were interleaved and all trials were presented in a fully random manner. In total, this protocol had a duration of 2320 sec.

#### Visual receptive field and tuning

To extract time-varying somatic calcium signals, we used the Suite2p toolbox (*55*). Neuropil contamination was corrected by subtracting the fluorescent signal from a surrounding ring F_Surround_(t) from somatic fluorescence: F(t) = F_Soma_(t) - U*F_Surround_(t), where U was estimated by the Suite2p deconvolution algorithm. Neuropil-corrected fluorescence signals F(t) where then converted in z-scores by subtracting from each trace the mean value and dividing by the standard deviation of F(t) over the samples contained in the time preceding stimulus onset in each trial (pooling across all trials). All data analysis of 2-photon single neuron responses was carried out with custom-made MATLAB software. Only neurons significantly responsive over at least one grating or sparse noise condition were included in the analyses of fig. 4. Responsiveness to each condition was assessed following the same procedure adopted in (*56*). The average of the z-scored calcium trace following stimulus onset (within the time window highlighted in fig. S8) was used to construct tuning curves and receptive field maps shown in Fig. 4C-D as well as S8. Preferred direction for each neuron was defined as the peak of the z-scored tuning curve. Preferred direction was defined in a similar way, after wrapping around direction from 0 to 360 deg to 0 180 deg to lump together conditions with the same grating orientation (irrespective of the direction of drift). To quantify orientation and direction selectivity OSI (Orientation Selectivity Index) and DSI (Direction Selectivity Index) were computed as defined in (*57*): OSI = (R_pref_ - R_ortho_)/(R_pref_ + R_ortho_), and DSI = (R_pref_ - R_opposite_)/(R_pref_ + R_opposite_), where R_pref_ is the response of the neuron to the preferred direction, R_ortho_ is the response to the orthogonal direction, relative to the preferred one (i.e., R_ortho_ = R_pref_ + n/2), and R_opposite_ is the response to the opposite direction, relative to the preferred one (i.e., R_opposite_ = R_pref_ + n). Receptive field (RF) location was estimated by performing a Gaussian fit of the combined ON/OFF receptive field map (obtained by averaging the absolute value of the ON and OFF fields). Pseudocolumns were defined by binning FOVs (Fields Of View) in up to 4 equi-sized spatial bins perpendicular to the manually annotated Cx-Ht boundary. Only pseudocolumns containing responsive neurons in both structures were used for the analysis described in Fig. 4D and S8C.

---

**Figure S1.**
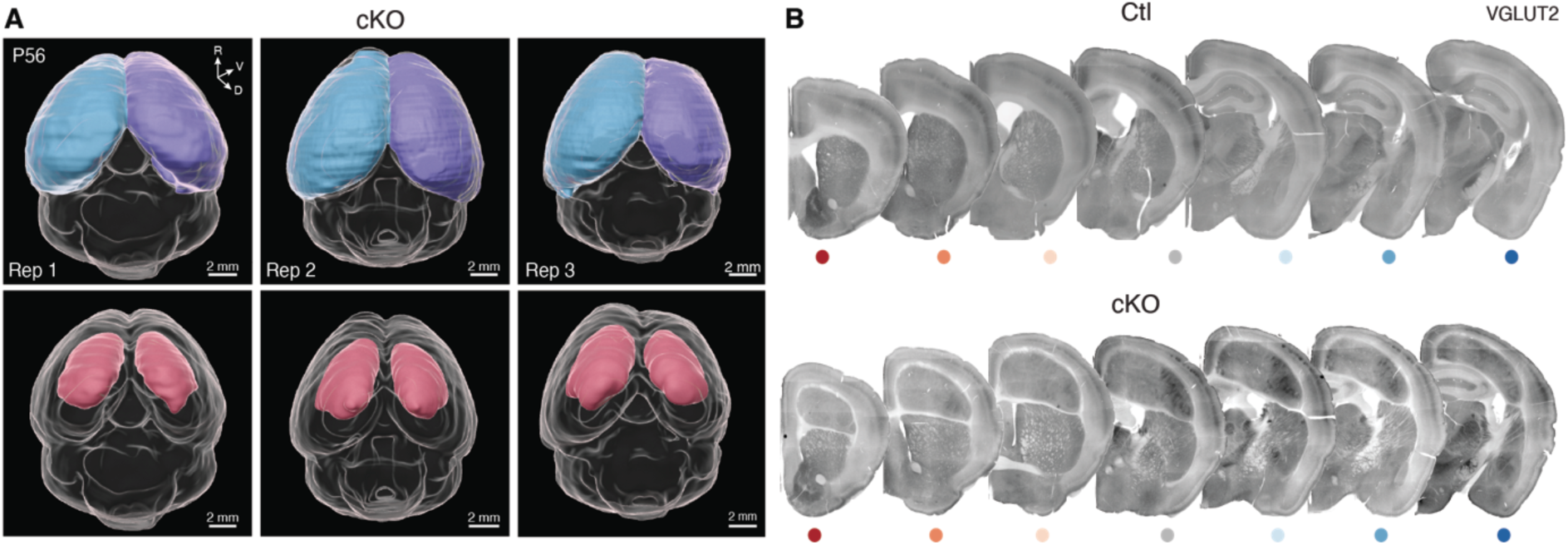
*Emll* dorsal pallium conditional deletion leads to a large subcortical band heterotopia. **(A)** 3D segmentation of cortical hemispheres (top) and bilateral Ht (bottom) in three adult cKO cleared brains. **(B)** Brain serial coronal sections of Ctl (top) and cKO (bottom) adult mouse brains stained for VGLUT2. Color-coded dots relate to the section rostro(red)-caudal(blue) level. Rep: replicates, R: rostral, D: dorsal, V: ventral.

**Figure S2.**
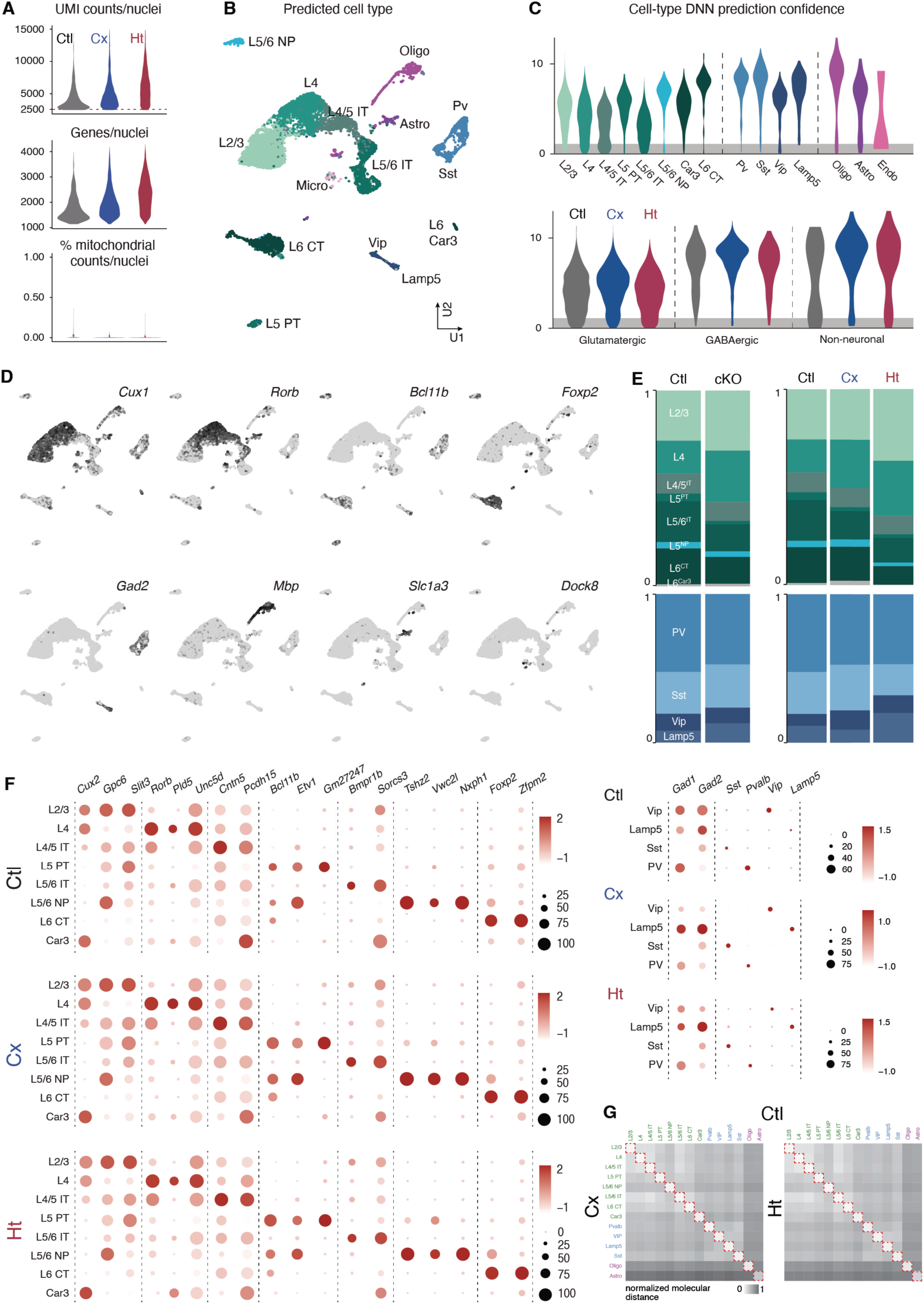
Diversity and molecular identities are robustly conserved in Ht. **(A)** Quality control features of Ctl, Cx and Ht sequenced nuclei. From top to bottom and per cell: detected number of RNA molecules, detected genes and mitochondrial counts. **(B)** UMAP representation of the whole snRNAseq dataset color-coded by DNN predicted cell-type (see Methods). **(C)** DNN prediction confidence by cell-type (top) and by sample cell-class (bottom). **(D)** Expression plots of cell-type specific glutamatergic, GABAergic and glial markers. **(E)** (Left) Ctl and cKO overall proportions of glutamatergic and GABAergic cell-types. (Right) Glutamatergic and GABAergic cell-type proportions by sample: Ctl, Cx and Ht. **(F)** Cell-type specific gene expression analysis (expression level and population proportion) of glutamatergic (left) and GABAergic (right) markers in Ctl, Cx and Ht types (top to bottom). **(G)** Molecular distance heatmap between Cx (left) and Ht (right) to Ctl cell-types. NP: near-projecting, IT: intra-telencephalic, CT: cortico-thalamic, PT: pyramidal tract, DNN: deep neuronal network.

**Figure S3.**
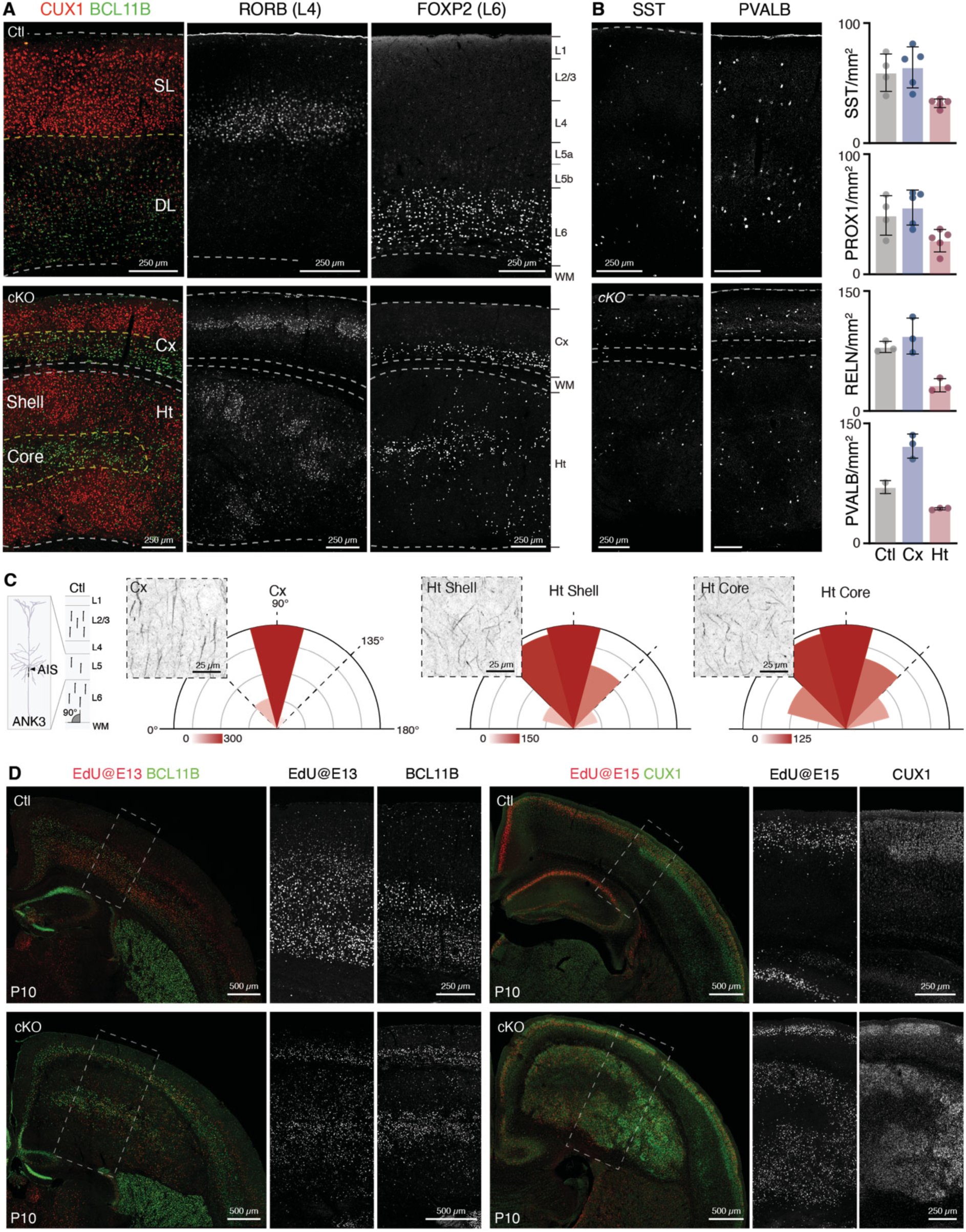
Ht neuronal identities and birthdate match despite their ectopic position. **(A)** Coronal section of Ctl (top) and cKO (bottom) stained for CUXl (layer 2/3), BCLllB (layer 5b, layer 6), RORB (layer 4) and FOXP2 (layer 6). Yellow-dotted line represents the limit between superficial and deep layer neurons in Ctl and between the core and the shell in cKO. **(B)** (Left) SST and PVALB staining in coronal Ctl (top) and cKO (bottom) brain sections. (Right) Density quantification of four GABAergic types in Ctl, Cx and Ht. **(C)** (Left) Schematic representation of axon initial segment staining and orientation in pyramidal neurons within a cortical column. (Right) Coronal images of Cx, Ht shell and core stained with ANK3 together with number of quantified AIS per orientation angle (see Methods) to the white matter. **(D)** Coronal brain sections of Ctl (top) and cKO (bottom) stained for EdU pulse at El3.5 and BCLllB (left) or EdU pulse at El5.5 and CUXl. SL: superficial layers, DL: deep layers, AIS: axon initial segment, EdU: 5’-ethynyl-2’-deoxuridine.

**Figure S4.**
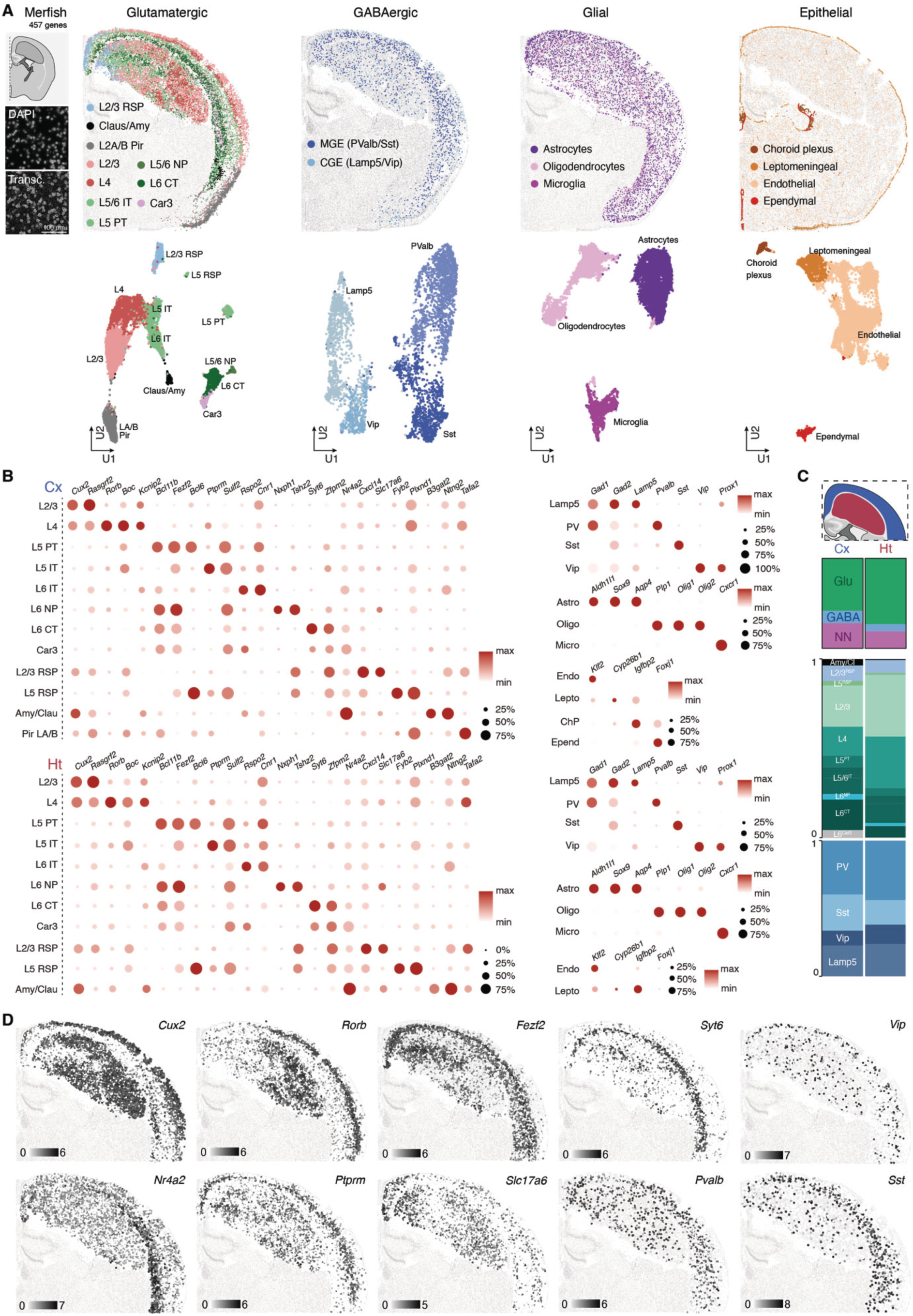
Spatial transcriptomics confirms cell type conservation in Ht. **(A)** (Left) Schematic representation of MERFISH *cKO* processed brain section. High magnification of DAPI nuclear staining along all the detected mRNA transcripts in the same field of view. (Right) Spatial distribution of glutamatergic, GABAergic, glial and epithelial cell-types over the processed brain section and UMAP representation of the different cell subsets color-coded by cell-type. **(B)** Cell-type specific gene expression analysis (expression level and population proportion) of glutamatergic (left) and GABAergic, glial and epithelial (right) markers in Cx and Ht types. **(C)** Cell-classes (top) and glutamatergic, GABAergic cell-type proportions (bottom) in Cx (blue) and Ht (red) segmented areas. **(D)** Spatial representation of **cross-normalized values of gene expression for** cell-type specific markers over Cx and Ht. Transc: transcripts, RSP: retrosplenial, Claus: claustrum, Amy: amygdala, Pir: piriform, NP: near-projecting, CT: cortico-thalamic, PT: pyramidal tract, Glu: glutamatergic, GABA: GABAergic, NN: non-neuronal.

**Figure S5.**
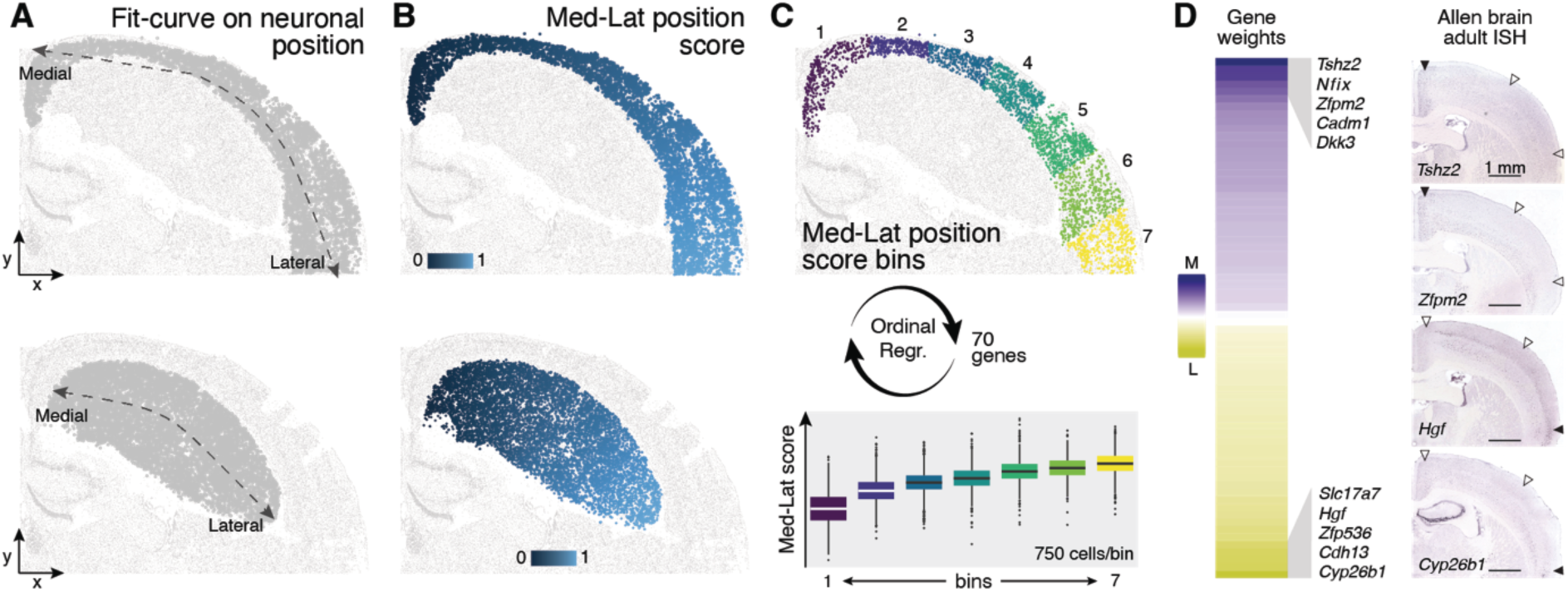
Ordinal regression model (ORM) used to predict cortical mediolateral molecular score. **(A)** Mediolateral position of Cx (top) and Ht (bottom) cells using with fit-curve function (see Methods) and **(B)** determination of single-cell mediolateral position score. **(C)** (Top) Representation of Cx mediolateral bins used to build the mediolateral ORM and (bottom) mediolateral molecular score boxplots of the ORM training cells. **(D)** (Left) ORM determined gene weights for the 70 model core genes (see Table 2) and (right) Allen Brain Institute ISH images of some selected candidate’s expression in Ctl brains. Med: medial, Lat: lateral, Regr: regression.

**Figure S6.**
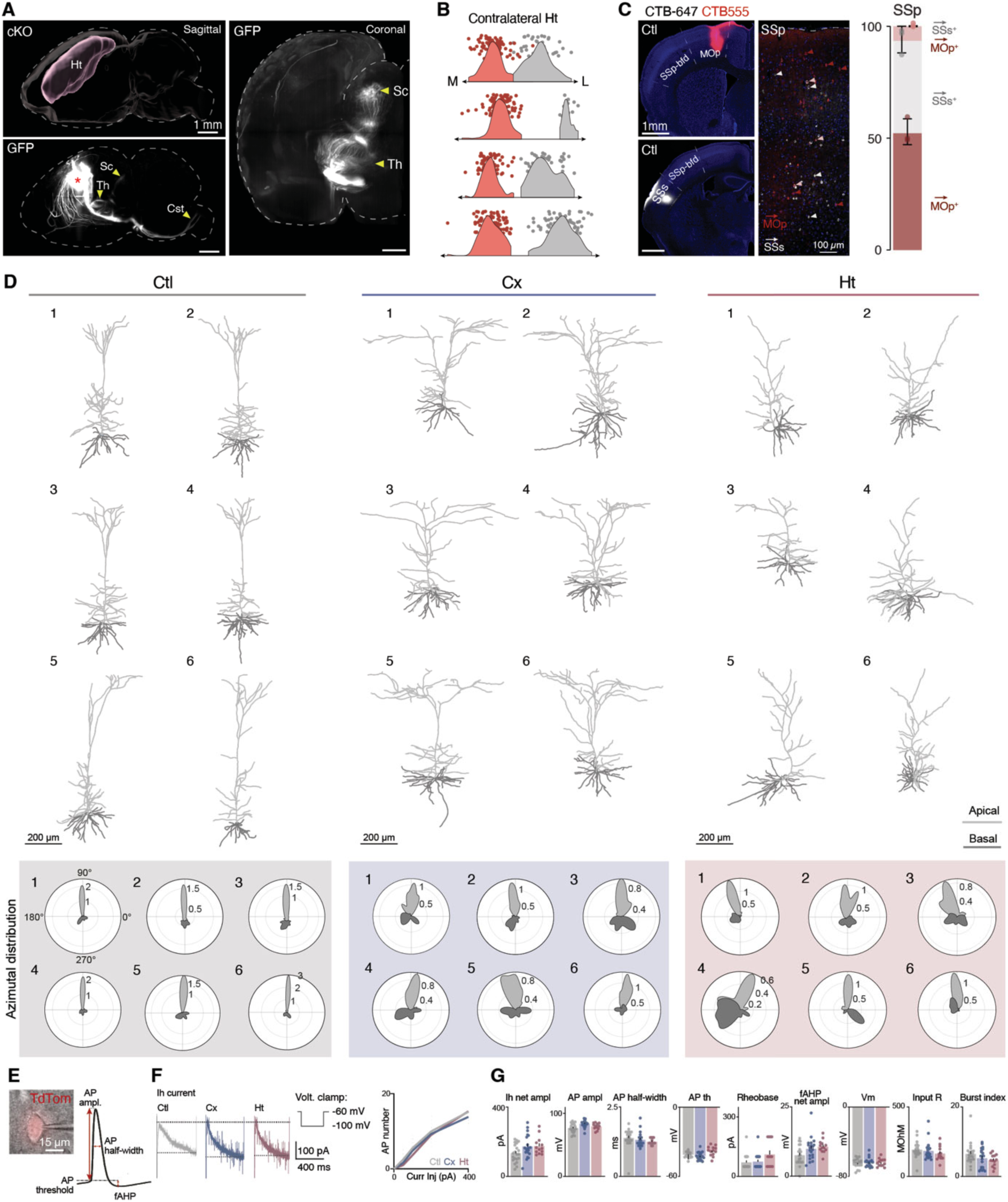
The output, morphology and electrophysiological characteristics of Ht neurons are preserved. **(A)** (Left) cKO cleared brain sagittal view of Ht segmentation (top) and GFP^+^ Ht anterograde projections (bottom). (Right) cKO cleared brain coronal digital section of GFP+ Ht anterograde projections to the thalamus and the superior colliculus. **(B)** Individual mediolateral distribution of medial and laterally retrogradely traced CC projecting neurons. **(C)** (Left) Ctl MOp (top), Ctl SSs (bottom) CTB injection sites and retrogradely labeled cells in an SSp column. Arrowheads point at MOp (red), SSs (light-gray) and dual (pink) projecting neurons. (Right) Cell-proportion of the three projection types in Ctl SSp. **(D)** (Top) Ctl, Cx and Ht L5b neurons morphology reconstruction (see Methods) displaying apical (light-gray) and basal (dark-gray) dendrites. (Bottom) Ctl, Cx and Ht single-neurons azimutal distribution of apical and basal dendrites density. Ctl neuron l, Cx neuron 6 and Ht neuron l are also displayed in fig. 2D **(E)** (Left) TdTom^+^ (CST projecting) patch-clamped neuron. (Right) Schematic representation of an action potential trend and its related measurements. **(F)** Ctl, Cx and Ht L5b neurons presentative Ih current (left) and excitability (right) curves. **(G)** Raw data of Ctl, Cx and Ht CST projecting neurons patch-clamp electrophysiological recordings (fig. 2E). GFP: green fluorescent protein, Sc: superior colliculus, Th: thalamus, MOp: primary motor cortex, SSp-bf: primary somatosensory cortex barrel-field, SSs: secondary somatosensory cortex, AP: action potential, Curr: current, Inj: injection, Volt: voltage, Ampl: amplitude.

**Figure S7.**
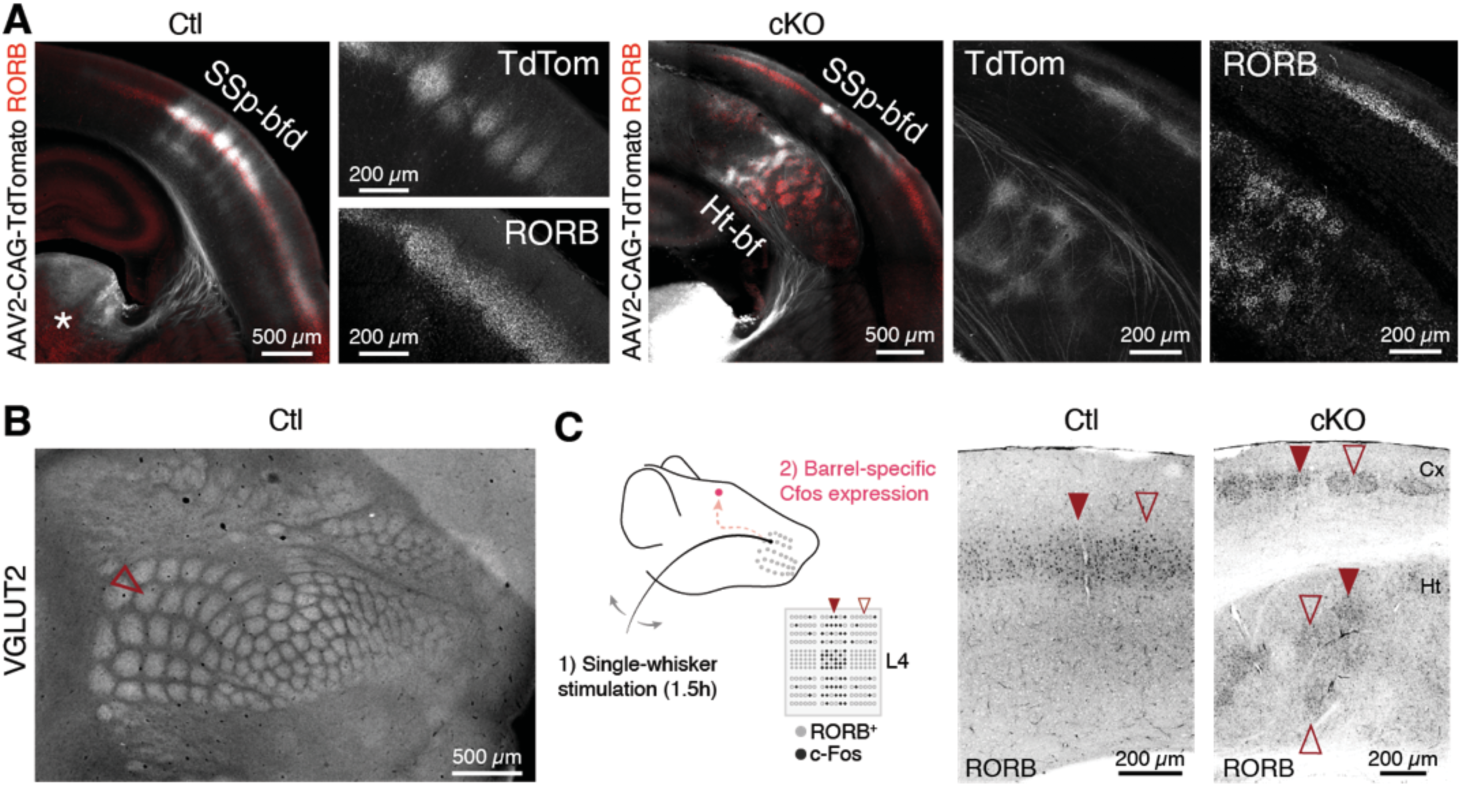
Thalamocortical terminals invade RORB^+^ barrel-like structures in the Ht. **(A)** Photomicrographs of thalamo-cortical projections anterograde tracing in Ctl (left) and cKO (right) animals with RORB immunolabeling. Low-magnification cKO image corresponds to fig. 3A but with RORB. **(B)** VGLUT2 labeling of thalamic terminals on Ctl cortex flattened preparations. Empty arrowhead points at the Bl barrel. **(C) (**Left) Schematic illustration of the single-whisker stimulation paradigm. (Right) RORB-expressing neurons corresponding to c-Fos^+^ neurons in fig. 3D. Full arrowheads highlight the active barrel and empty ones, the inactive ones. SSp-bfd: primary somatosensory cortex barrel-field.

**Figure S8.**
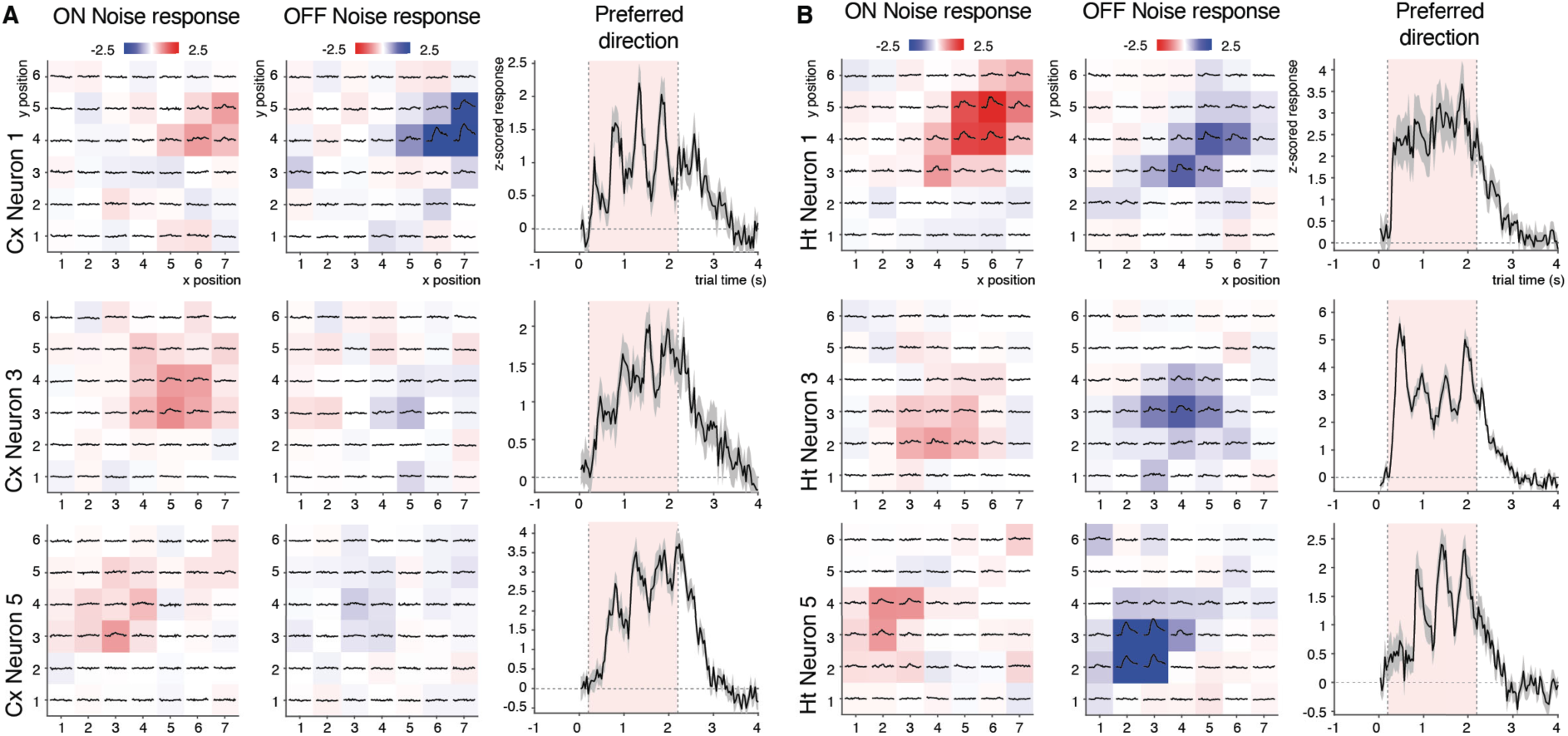
Ht neurons below VISp show similar receptive field distribution to cortical counterparts. **(A)** Cx and **(B)** Ht neuronal ON and OFF responses for each spatial location corresponding to neurons l, 3 and 5 (fig. 4A). Black curves in each spatial location represent the average response time course of a neuron when the stimulus was presented at that location. (Right) Average response time course of each neuron when presented with the drifting grating of preferred orientation.

**Figure S9.**
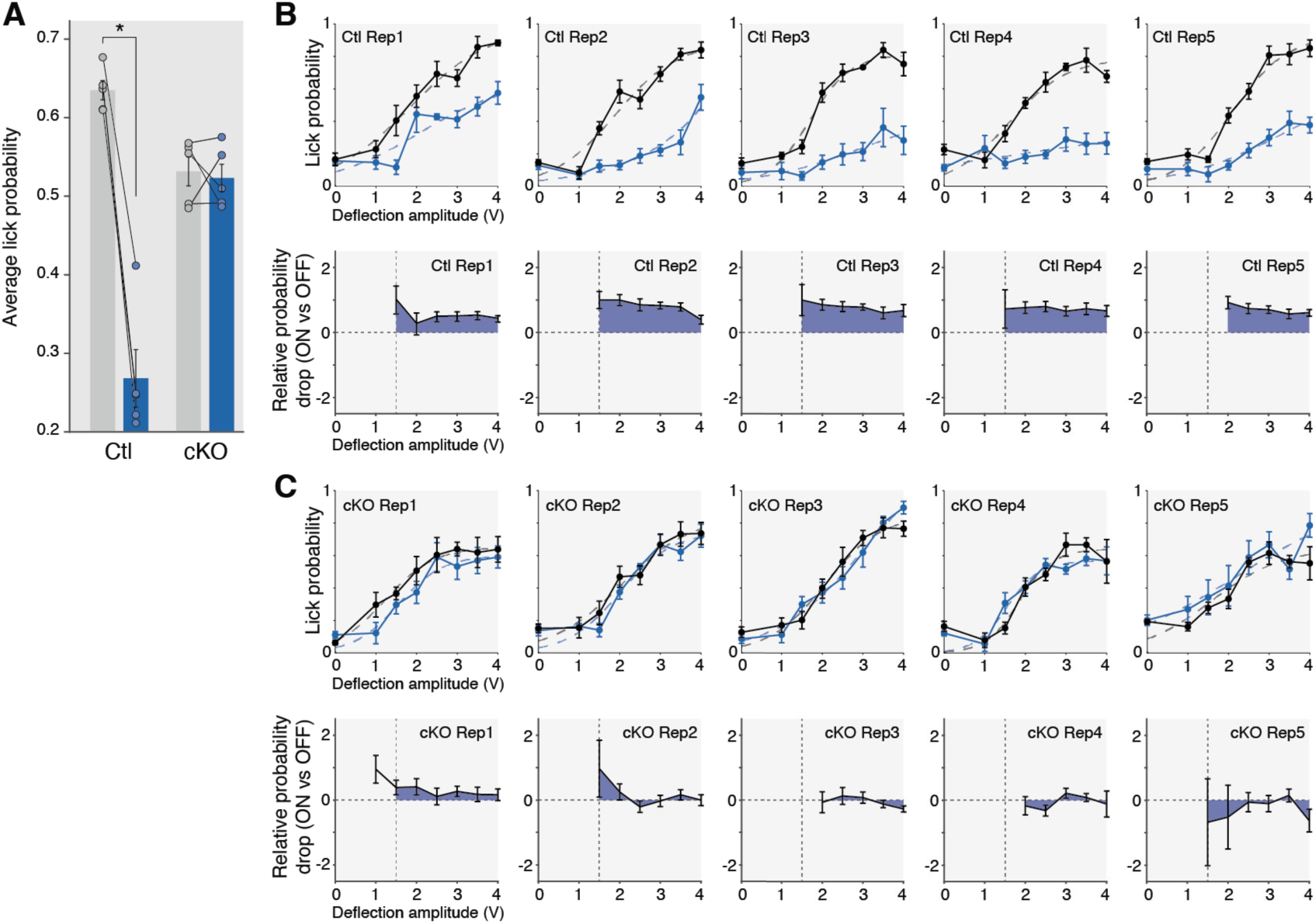
Ht neurons replace inhibited cortical functions. **(A)** Average licking probability drop per animal between ON and OFF conditions for Ctl (paired t-test, *P* = 0.000l)and cKO (paired t-test, *P* = 0.74) animals. **(B)** (Top) Licking probability of individual Ctl and **(C)** cKO mice under normal conditions (OFF - black) and blue light stimulation (ON - blue) for different whisker deflection intensities. (Bottom) Individual relative licking probability loss between OFF and ON conditions. Rep: replicate.

